# Visualizing structure-mediated interactions in supercoiled DNA molecules

**DOI:** 10.1101/257949

**Authors:** Shane Scott, Zhi Ming Xu, Fedor Kouzine, Daniel J. Berard, Cynthia Shaheen, Barbara Gravel, Laura Saunders, Alexander Hofkirchner, Catherine Leroux, Jill Laurin, David Levens, Craig J. Benham, Sabrina R. Leslie

**Affiliations:** Department of Physics, McGill University, Montreal, Quebec, Canada H3A 2T8; Center for Cancer Research, National Cancer Institute, Bethesda, Maryland 20892; Genome Center, University of California Davis, Davis, California 95616

## Abstract

We directly visualize the topology-mediated interactions between an unwinding site on a supercoiled DNA plasmid and a specific probe molecule designed to bind to this site, as a function of DNA supercoiling and temperature. The visualization relies on containing the DNA molecules within an enclosed array of glass nanopits using the Convex Lens-induced Confinement (CLiC) imaging method. This method traps molecules within the focal plane while excluding signal from out-of-focus probes. Simultaneously, the molecules can freely diffuse within the nanopits, allowing for accurate measurements of exchange rates, unlike other methods which could introduce an artifactual bias in measurements of binding kinetics. We demonstrate that the plasmid’s structure influences the binding of the fluorescent probes to the unwinding site through the presence, or lack, of other secondary structures. With this method, we observe an increase in the binding rate of the fluorescent probe to the unwinding site with increasing temperature and negative supercoiling. This increase in binding is consistent with the results of our numerical simulations of the probability of site-unwinding. The temperature dependence of the binding rate has allowed us to distinguish the effects of competing higher order DNA structures, such as Z-DNA, in modulating local site-unwinding, and therefore binding.

## INTRODUCTION

Interactions between nucleotides and other molecules are an important part of cellular function: when either DNA replication or transcription are incorrectly performed, their parent cell may suffer. Other processes, such as those during homologous recombination, require the binding of freely-diffusing single-stranded oligonucleotides (oligos) to an unwound region on double-stranded DNA (1, 2). R-loops, formed by RNA oligonucleotides bound to DNA, are now known to possess a role in mitochondrial and bacterial replication, eukaryotic transcription, and cancer (6). Additionally, new biotechnologies such as CRISPR-Cas9 use single guide RNA (sgRNA) oligos to bind to specific regions on target DNA molecules (3, 4). Deviations from the expected interaction can have severe consequences: in the case of DNA replication, permanent errors in the genetic code could lead to mutations in the genome, causing diseases such as cancer (5).

One of the key elements for many of these processes to occur is the single-stranded separation of DNA, or unwinding. Opening double-stranded DNA makes it accessible to replication and transcription machinery, as well as allowing access to DNA or RNA oligos. Illuminating DNA unwinding and subsequent binding is important for understanding these processes, and could serve to improve nucleotide biotechnology and gene editing techniques such as CRISPR-Cas9.

Several variables are known to drive DNA unwinding, such as temperature, DNA-protein interactions, and DNA supercoiling. DNA supercoiling is prevalent in physiological systems, both prokaryotic and eukaryotic (7, 8, 9, 10, 11). Here, the cells bind enzymes called topoisomerases to DNA, directly affecting its supercoiled state. Many other DNA-binding proteins are known to affect the supercoiled state of DNA. RNA polymerase, for example, is known to dynamically change the supercoiled state of DNA, sufficiently so that it may cause a specific DNA region to unwind (12). The dynamics of supercoil-induced structural transitions in DNA are important to a wide range of cellular processes, such as chromosome reading during replication and transcription.

Proteins, RNA, or higher-order complexes can bind to and dynamically change the supercoiled state of DNA, which can influence the state of genes through unwinding (11, 12). Torsionally induced unwinding is also prevalent during homologous recombination, a process that has been proposed to be initiated by specific, supercoil-sensitive regions called D-loops. These areas possess oligonucleotides annealed to one strand in unwound regions, which are thought to be significantly influenced by supercoiling (1, 2), an open question for investigation. In cells, DNA unwinding fluctuations can also explore different structural states, which is potentially important for gene expression.

### DNA Supercoiling

To build a model of how DNA structure mediates site-unwinding and binding, we introduce supercoiling as the sum of the over- or under-twisting (*Tw*) present in the DNA helix, and the writhe (*Wr*), which describes the large-scale geometry of the molecule — that is, the crossing of one part of the DNA helix over another (13, 14). DNA supercoiling can drive local structural transitions such as from B- to Z-form or duplex unwinding at susceptible sites (Fig. 1) (1, 15, 16, 17). These alternate structures can in turn serve as initiation sites for processes that require template DNA insertion, such as DNA replication and repair (17, 18, 19).

**Figure 1.**
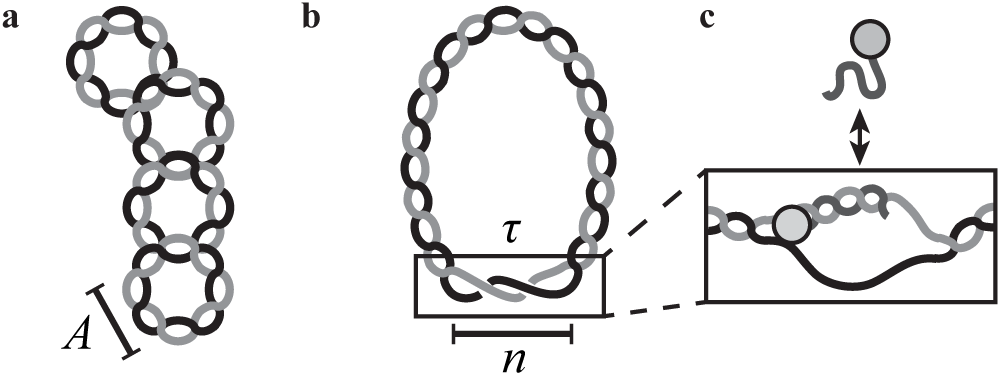
DNA supercoiling promotes unwinding. **a)** Supercoiled plasmid DNA experiences stress due to either over- or under-twisting of the DNA double helix. The torsional stress caused by this can force the DNA double helix to cross over itself, or affect the average number of base pairs (bp) per complete turn of the double helix, *A* = 10.4 bp/turn. **b)** Sufficiently high stress allows for the structural transitions along the molecule, such as DNA unwinding. Here, *n* bases are single-stranded, or unwound, and wrap about each other by a certain number of radians per base, *τ*. **c)** DNA probe molecules bind to target sites only after they become unwound, allowing for visualization of unwinding.

Supercoiling can drive site-unwinding by introducing torsional stress. This is quantified by the superhelical density, 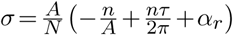. Here, *A* is the average number of base pairs (bp) per turn in an unstressed DNA molecule (10.4 bp per right-handed turn), *N* is the total number of base pairs, *n* is the number of unwound base pairs, and *τ* is the torsional deformation rate of the open region and reflects how the unpaired strands twist around each other. Further, *α*_*r*_ is the residual supercoiling, which includes the other structural effects, such as *Wr* and torsional deformations of the B-form regions (17, 20). A negative value of *σ* corresponds to underwound DNA, and a positive value corresponds to overwound DNA.

Despite the importance of supercoil-induced unwinding dynamics in DNA, microscopy techniques have been limited in visualizing and investigating these changes without perturbing the system mechanically in ways that restrict the accessible conformational states of the molecules and thereby affect the results (21). Here, we present a single-molecule microscopy technique that allows us to probe DNA unwinding as a function of supercoiling and temperature without artificially perturbing the DNA structure.

## MATERIALS AND METHODS

While many techniques have been developed to observe how DNA tension and torsional strain regulate biomolecular processes, they typically require artificially constraining DNA structure (e.g. with surface-tethers). These modifications are essential to the techniques: without them, the molecules would diffuse out of view within milliseconds. Unfortunately, the modifications typically disturb the DNA secondary structure and limit the accessible conformations, thus affecting measurements sensitive to DNA structure (22, 23, 24). Gels are also frequently used to constrain and study supercoiling in DNA, but provide only a static, population-averaged picture and lack the ability to follow dynamics and kinetic events in real-time (25).

Here, we demonstrate a direct, single-molecule approach to studying topology-mediated dynamics and interaction kinetics of biomolecules in solution while simultaneously leaving them free to explore all possible structural conformations. Our method combines the Convex Lens-induced Confinement (CLiC) technique (26, 27, 28) with a nanofabricated array of pits that are etched in the bottom surface of a deformable glass flow-cell. In our experiments, we load and trap DNA oligo-probes and plasmids in individual pits so they are free to diffuse, fluctuate, and interact. Due to their simplicity, robust nature, and availability, we chose to use DNA oligos instead of RNA. The pits are significantly larger than the molecules, leaving the latter free to explore all available conformations; they are not affixed, for instance, by chemical tethers or beads (Fig 2).

**Figure 2.**
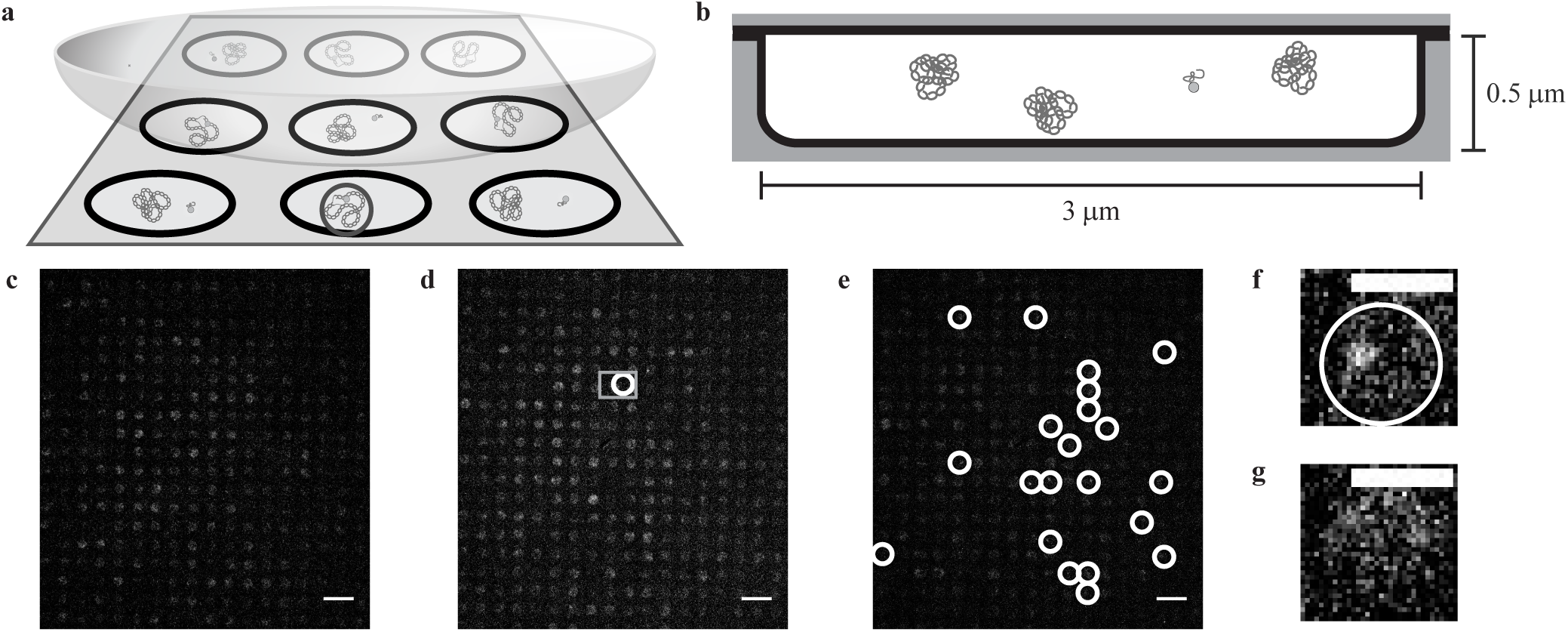
CLiC nanopit microscopy and binding events. **a)** Schematic of the CLiC method confining plasmids and labeled oligo-probes to pits made using micro/nano lithography techniques (28). The circle in the front-center pit demonstrates a pit containing a bound probe-plasmid complex. **b)** Schematic of plasmids and a freely diffusing probe confined to a circular pit of 3-*µ*m diameter and 0.5-*µ*m depth. Each pit contains ∼24 plasmids and a single labeled probe. **c)** Fluorescence image of mixture of probes and pUC19 plasmids with mean superhelical density < *σ* >= 0 and standard deviation *sσ* = 0.03 confined to pits. **d)** Fluorescence image of mixture of probes and pUC19 plasmids with superhelical density < *σ* >= 0.050, *sσ* = 0.07 confined to pits. The gray square denotes the two pits containing single probes in f and g. **e)** Fluorescence image of mixture of probes and pUC19 plasmids with < *σ* >= *-*0.127, *sσ* = 0.06 confined to pits. White circles show binding events. **f)** Pit containing a bound probe-plasmid complex from the data set shown in d. **g)** Pit containing a freely diffusing probe from the data set shown in d. Scale bars in c)-e) denote 10 *µ*m, while those in f) and g) are 3 *µ*m. Images in c)-g) had their background subtracted using a rolling ball algorithm with a radius of 50 pixels, and a Gaussian-blur filter with radius 0.5 pixels was applied to each.

The 500 nm-deep pits are within the focal depth, so once sealed in pits, molecules are confined to the focal plane and are thus always in view. The nanopits are organized into an array, which is sealed by the CLiC method during imaging. This provides multiple independent reaction chambers in which the molecules can be observed under a large dynamic range of reagent concentrations and observation timescales. Each chamber can be emptied and replenished with a new sample of molecules from bulk solution by lifting and lowering the CLiC lens. Using precision-fabricated glass pits enables quantitative, contaminant-free, and high-quality visualization of molecules in solution. Previous studies involving individual reaction chambers have used either resealable pits with at least one polydimethylsiloxane (PDMS) surface (29, 30, 31), or permanently sealed fused silica pits (31). PDMS can be susceptible to absorption of reagents and exchange of oxygen with the surrounding environment, as well as swelling, and is characterized by a larger autofluorescence than glass. Our technique allows for sample replenishment of pits (and hence serial experiments to be performed using the same device, with high statistics), prevention of oxygen exchange (and hence suppression of photobleaching, with extended observation periods), suppression of autofluorescence, and precision characterization of the fabricated glass nanopits.

To visualize supercoil-induced DNA unwinding, we developed an assay that uses probe-binding to target specific regions on highly underwound plasmids. The DNA was sufficiently underwound to drive local strand-separation, making the specific regions accessible to the probes (Fig. 1c). We used pUC19, a model plasmid constructed using sequences from the plasmid pBR322, as it is well studied and widely used as a cloning vector (32, 33). pUC19 possesses two supercoil-induced unwinding regions called Site 1 and Site 2 (15, 34). Site 1 is associated with the origin of replication and has the largest probability of unwinding at most superhelical densities. Site 2 is associated with an ampicillin resistance gene and only unwinds under large torsional stress (15, 35).

The plasmids were isolated from DH5*α E. coli* cells grown in our lab. Topoisomers were produced by treating these plasmids with a topoisomerase 1B from calf thymus in various amounts of ethidium bromide. For an in-depth discussion of our plasmid growth, purification procedures, and enzymatic reactions, please refer to SI.

For this work, we designed a series of single-stranded, fluorescently-labeled DNA oligo-probes to be complementary to one side of Site 1 in pUC19 (see SI), either at the edge or the center of the predicted unwinding region. The logic behind our assay is that, should Site 1 unwind and become single-stranded, we would expect a probe to bind to its complementary sequence, forming a probe-plasmid complex. We conjectured that a sufficiently long probe, once bound, would remain so during the experimental timescale (refer to SI). Consequently, the locally denatured “DNA bubble” would be maintained for clear visualization.

By confining plasmids and probes to a pit array using a glass lid, we can visualize the interactions between the plasmid unwinding site and probes as a function of *σ* and temperature (Fig. 2a). Each pit is approximately 3 *µ*m in diameter and 500 nm in height. The nanoscale smoothness of the glass surfaces allows the pits to be sealed, preventing molecules as small as our probes (*∼*1 nm in size) from escaping during observation. Furthermore, the sealed geometry dramatically reduces photobleaching by suppressing oxygen exchange, allowing for extended per-molecule observation. The exclusion of out-of-focus fluorophores by the CLiC method enables sensitive detection of single labels. The tight seal and well-defined, impermeable geometry are enabled by the use of glass as a confining surface. For an in-depth discussion of the tools, experiment preparation, and microscopy technique, refer to SI.

We distinguish between fluorescence images of diffusing probe-plasmid complexes and freely diffusing probes by their differences in diffusivity, size, and intensity distribution. Using a finite imaging exposure time (e.g. 50 ms/frame), the fluorescence from unbound, diffusing probes uniformly illuminates a pit. By contrast, the bound probe/plasmid complex appears as a bright, diffraction-limited spot. Tracking of the confined molecules allows us to probe the state of the interaction as it progresses, with high statistics. For a more in-depth discussion of our binding detection algorithm, refer to SI.

The interaction between the unwound plasmid *U* and probe *P* can be considered as a reaction between two reactants to form a product *UP*. As we found that the probe-plasmid complex is very stable at experimental temperatures and time-scales, we modeled the interaction as a second-order reaction where *U* +*P*→*UP*. This model assumes that the opening and closing of unwinding sites occurs on average at timescales longer than one of our experiments. Further discussion on the validity of this assumption can be found in the Results section.

As we cannot directly measure the concentration of unwound plasmids [*U*] but we can count the number of bound complexes [*UP*]=[*U*]_0_ − [*U*] with [*U*]_0_ being the initial unwound plasmid concentration, we modified the second-order reaction equation. We also assumed that the probe molecules are in excess to the bound complexes for all experimental conditions such that [*P*]≈[*P*]_0_. This assumption was determined experimentally to be valid, as will be discussed in the Results section. Taking these conditions under consideration, we obtain

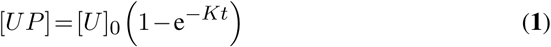

where *K* = *k* ([*P*]_0_ -[*U*]_0_), *k* is the true rate constant for this reaction, *t* is the time, and [*P*]_0_ is the initial concentration of probes.

## RESULTS

### Guiding predictions: theoretical treatment of unwinding

DNA typically exists as B-DNA, a right-handed double helix with an average number of bases per turn given by *A* = 10.4 bp/turn at physiological conditions. This structure is most stable when a DNA molecule is not under torsional stress (Fig. 3a). At low supercoiling levels, supercoiling is typically partitioned between twist and writhe (Fig. 3b, c). Once enough supercoiling is added, the energy may also be partitioned to create other non-B-DNA conformations, such as Z-DNA (a left-handed double helix), cruciforms, or unwound strands, with the latter shown in Fig. 3d.

**Figure 3.**
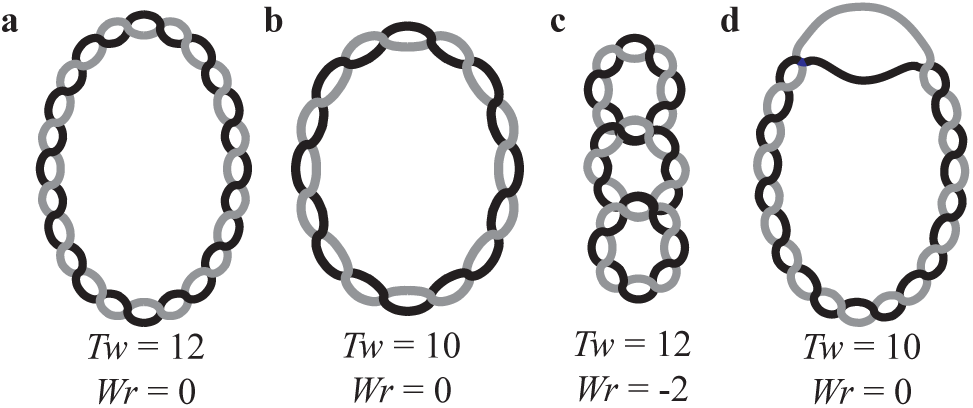
Partitioning of supercoiling. **a)** A completely relaxed plasmid molecule. There are 12 turns of the double helix, and, as the helix does not cross over itself, the twist (*T w*) is equal to 12 and writhe (*W r*) is equal to 0. **b)** The same plasmid molecule as in a), but supercoiled. The double helix has been untwisted by 2 turns but does not cross over itself, leading to *T w* = 10 and *W r* = 0. **c)** The supercoiled molecule in b can release some of the undertwisting stress energy by crossing the double helix over itself. Here, *T w* = 12 while *W r* = -2. **d)** Another means of relieving stress energy is through site unwinding, depicted here for the same plasmid as in b and c. Whether a plasmid partitions its energy into *T w, W r*, or other secondary structures, such as unwinding, is probabilistic for each base pair, and depends on its local energy landscape.

While we do not have a theoretical model of the site unwinding kinetics and dynamics, we do have an equilibrium model of strand-unwinding to inform our experiments. Here, we use the Stress-induced Duplex Destabilization (SIDD) model to predict unwinding, as well as related multi-state competitive transition models (e.g. DZCBtrans), all of which are grounded in statistical mechanics (17, 20, 35). These models predict that sufficient supercoiling can induce regions of alternate secondary structure, including unwound regions, cruciforms, and Z-DNA (see SI). Briefly, each base pair along a DNA molecule can exist in B-form, or as another secondary structure. Supercoiling can induce competition among the energetically favored secondary structures, which are analyzed in the DZCBtrans model; this model provides a comprehensive prediction of the secondary structure in a given DNA molecule at equilibrium as a function of *σ*, temperature, and ionic strength.

For a given molecule of DNA, we consider the transition to any non-B-DNA state *t*. For *n*_*t*_ base pairs to undergo this transition, there must be enough energy to nucleate it, plus additional energy 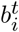 to change the state of every individual base pair *i*. Here we assume that the total number of possible DNA conformations is *m*. For the DNA molecule to exist in a state with multiple different DNA conformations, it must possess a free energy of

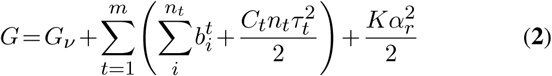

where *G*_*v*_ is the total nucleation energy for all the conformational transitions, *C*_*t*_ is the torsional stiffness coefficient for conformation *t*, and *K* is the superhelical energy parameter governing *α*_*r*_ (17). Energies of twisting for the alternative conformations are considered in the third term, while the energetics of DNA writhe are included in the fourth term. Both SIDD and the DZCBtrans model may be described using this formalism, with the former using *m* = 1 and the latter *m* = 3. Values of the parameters 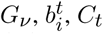, and *τ*_*t*_ must be known for each conformation included.

A partition function can be constructed and probabilities for the states of the molecule can be calculated. Here, the probability of the DNA molecule existing in a certain state decreases exponentially as the energy of the state increases. The most likely conformation of a given molecule of DNA is therefore the state that minimizes the free energy, *G*. From this information, the probability of transition of each base pair can be computed. For further information, refer to Zhabinskaya and Benham (17).

In applying this theory to the pUC19 plasmid system, we took into account the Gaussian-distributed topoisomer superhelical density with a mean of < *σ*> and standard deviation of *s*_*σ*_ = 0.007 of the experimental sample (see SI for the experimental determination of these values). Our calculations predict that the competition with Z-DNA formation strongly affects the occurrence of supercoil-induced unwinding. SI provides a discussion of the calculations and a comparison to the predicted unwinding behavior without the Z-DNA competition.

DZCBtrans predicts a strong thermal dependence of unwinding (Fig. 4). Briefly, the Z-DNA transition is predicted to be favored over strand-denaturation at low temperatures (see SI); this formation of Z-DNA is responsible for suppressing supercoil-induced unwinding of Site 1, even at highly underwound supercoiling. As the temperature is increased, more supercoil-induced unwinding is predicted to occur, while the presence of Z-DNA is predicted to decrease. (Note that the decrease in the number of unwound bases predicted by SIDD coincides with some supercoiling energy being used to unwind Site 2 on the plasmid.) DZCBtrans theory is consistent with our experimental observation of a strong dependence of oligo-probe site binding on temperature, and the interpretation that this requires site-unwinding; whereas SIDD theory is not.

**Figure 4.**
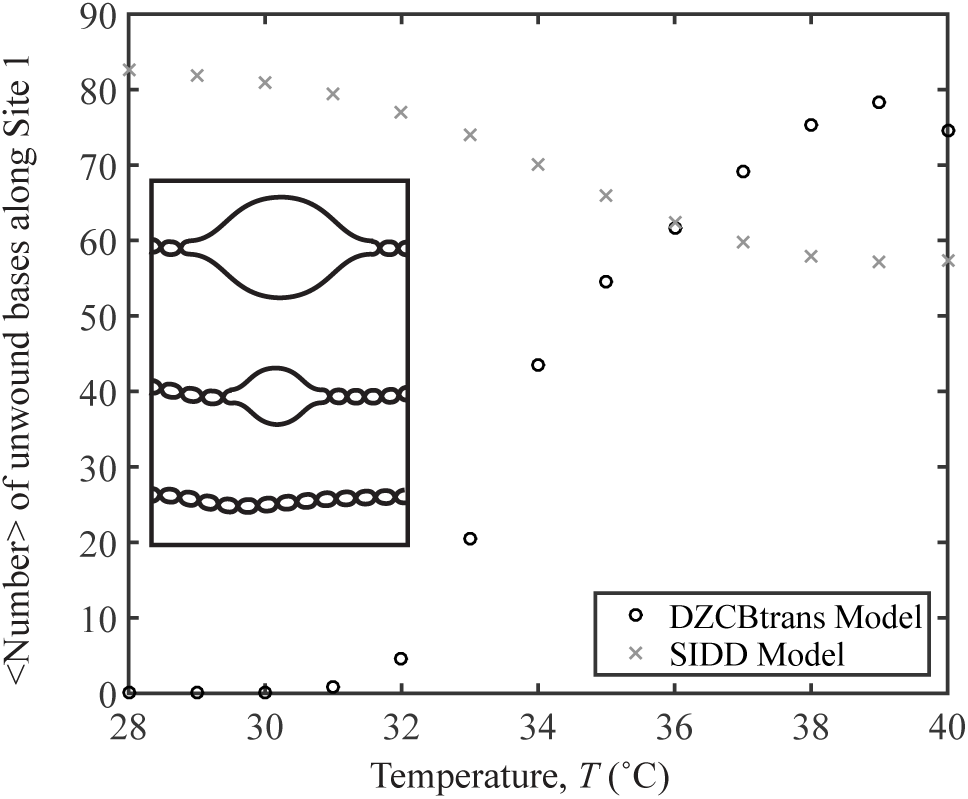
Statistical mechanics predictions of unwinding. DZCBtrans and SIDD profiles for mean superhelical density < *σ* >= -0.120 with standard deviation *sσ* = 0.007 as a function of temperature for the Site 1 region of pUC19. Inset: schematic of Site 1 unwinding.

### Experimental results: site binding by oligo-probes increases as a function of temperature

Fig. 5 demonstrates an increase in the interactions between the probe and the unwinding region with temperature. At each temperature, a separate sample was prepared with a plasmid superhelical density of < *σ*>= -0.108, each with *s*_*σ*_ = 0.007 (see SI for a discussion of topoisomer determination and microscopy procedure). Fig. 5a shows the time-evolution of the number of bound complexes, which is characterized by a greater rate at higher temperatures. Fits to the data were made using eq. **1**, and are presented here as curves. The fits and data both seem to approach a certain number of bound complexes at long timescales. As only one oligo may bind per plasmid and not all oligos have reacted, this indicates that the initial number of unwound regions on pUC19 are constant during the experimental time frame and validates our assumption that the opening and closing of the unwinding region is slow compared to the reaction. Additionally, the maximum number of bound complexes is just over 10 per 100 oligos, justifying the second assumption that [*P*]≈[*P*]_0_. The number of bound complexes in each video was measured using the counting algorithm described in SI, and the results per pit array were scaled per 100 probes.

**Figure 5.**
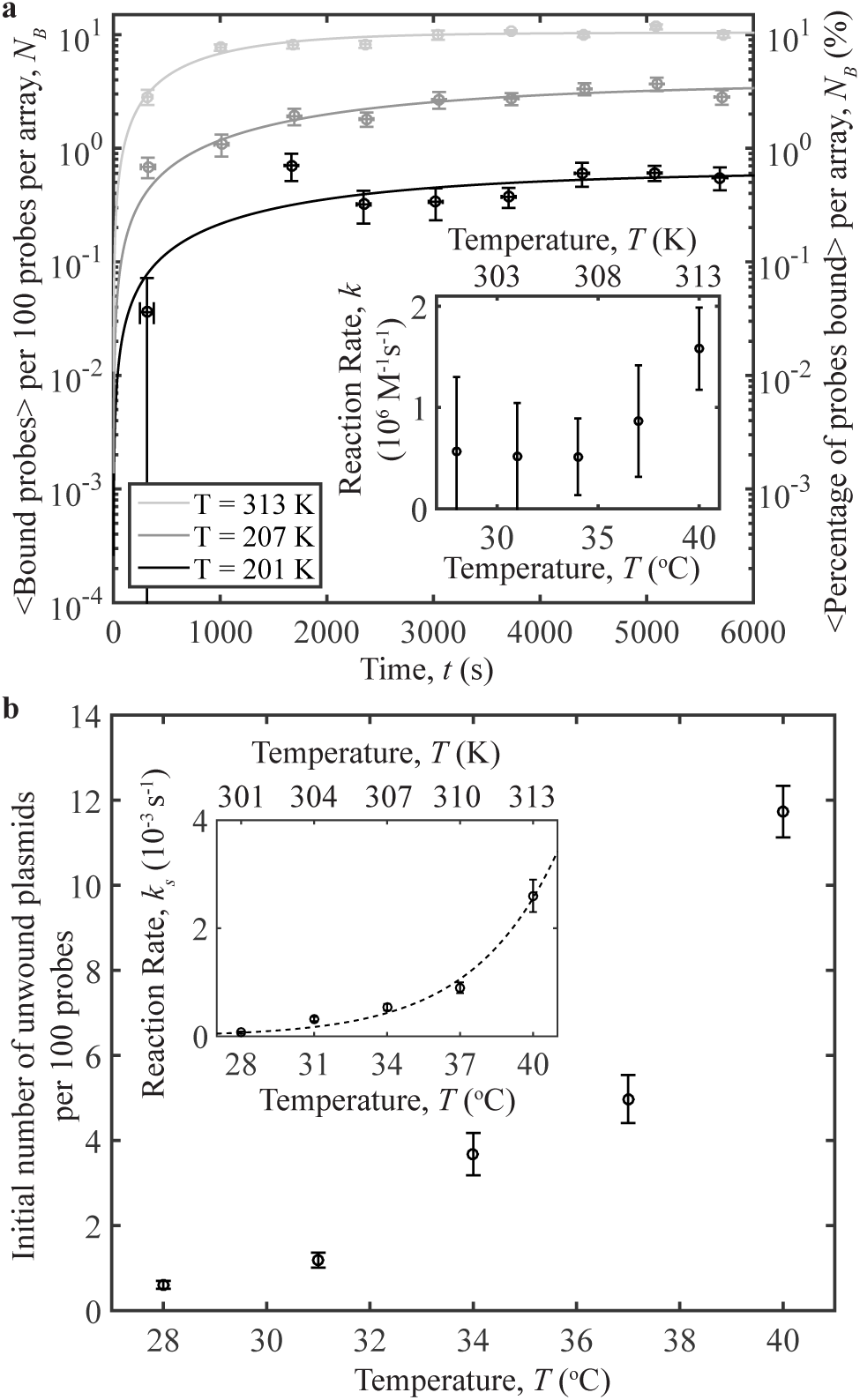
Probe-plasmid binding and temperature. **a)** The number of binding events per 100 probes averaged over 10 different videos vs. time, for three temperatures studied. Curves represent fits to the data. The plasmids have a superhelical density of < *σ* >= -0.108, *sσ* = 0.07. Inset: Reaction rates vs. temperature obtained from the data fits. **b)** The number of unwound plasmids obtained from the fits in a) per 100 probes vs. temperature. Inset: the short-time reaction rate, *ks*, vs. temperature for the data in part a. The reaction rates are fit to an exponential demonstrating Arrhenius behavior. Each data point was averaged over ∼3,300 probes, indicating high-throughput. Error bars in a) and the inset of b) denote the standard error of the mean of the binding for the 10 averaged videos on the y-axis, while those on the x-axis represent the standard error of the mean of the time. Error bars on the fit parameters represent the 95% confidence interval of the fit.

Results from the fits to the data were not obvious as two processes must occur in order for our technique to detect binding events: site-unwinding followed by probe binding. The increase in the number of pUC19 molecules with unwound regions per 100 probes vs. temperature is plotted in the inset of Fig. 5b. The observations are consistent with site-denaturation allowing binding of probes. The observed strong temperature dependence agrees with the predictions by DZCBtrans in contrast to that of SIDD theory (Fig. 4). These results illustrate the critical role of secondary structures, namely Z-DNA, in mediating site-specific interactions. See SI for experimental evidence of, and discussion on, the presence of secondary structures in pUC19.

Both the number of unwound plasmids and the overall reaction rate increases with temperature, as should be expected: higher temperatures increase diffusion leading to more encounters between reactants and thus a higher rate of reaction (inset of Fig. 5a), and provide more energy, promoting unwinding. While the predicted behavior of the reaction rate should be exponential, the large 95% confidence intervals on the fit parameters render a more exact determination of the thermal-dependence of the reaction difficult. To more accurately demonstrate the reaction’s thermal dependence, we calculated the short term reaction rates (*k*_*s*_) by approximating a linear fit to the short-time data. The measurements of *k*_*s*_ depend exponentially upon temperature, consistent with Arrhenius behavior (Fig. 5b inset). High statistics offered by the CLiC-nanopits visualization technique provide direct access to the weak topology-mediated probe-plasmid interactions, occurring over hundreds to thousands of seconds. For example, on average ∼3,300 probes were sampled for each data point in Fig. 5, indicating its value as a high throughput technology.

### Experimental results: oligo-probe binding to plasmid unwinding sites increases as a function of superhelical density

In a subsequent series of experiments, samples were prepared as before, but experiments were conducted using a series of <*σ*> values and a constant temperature of 37°C. Topoisomers with different < *σ*>, all with *s*_*σ*_ = 0.007 were prepared using topoisomerase IB and a range of ethidium bromide concentrations (described in detail in SI). A set of videos was acquired as before, and the number of bound complexes in each video was measured and scaled per 100 probes (Fig. 6a). Fits for each data set to eq. **1** are presented through curves. As superhelical tension is added to the system there is an increase in probe binding; this trend extends to less negatively supercoiled samples. Reaction rates are shown in the inset of Fig. 6a. These rates are constant, consistent with the idea that increasing supercoiling induces unwinding, not an increase in collisions between the reactants. In Fig. 6b, the number of unwound pUC19 molecules per 100 probes determined from the fits is shown vs. < *σ*>. The “short-time binding rates” were calculated as in the previous section; they are shown in the inset of Fig. 6b to increase with the level of negative supercoiling.

**Figure 6.**
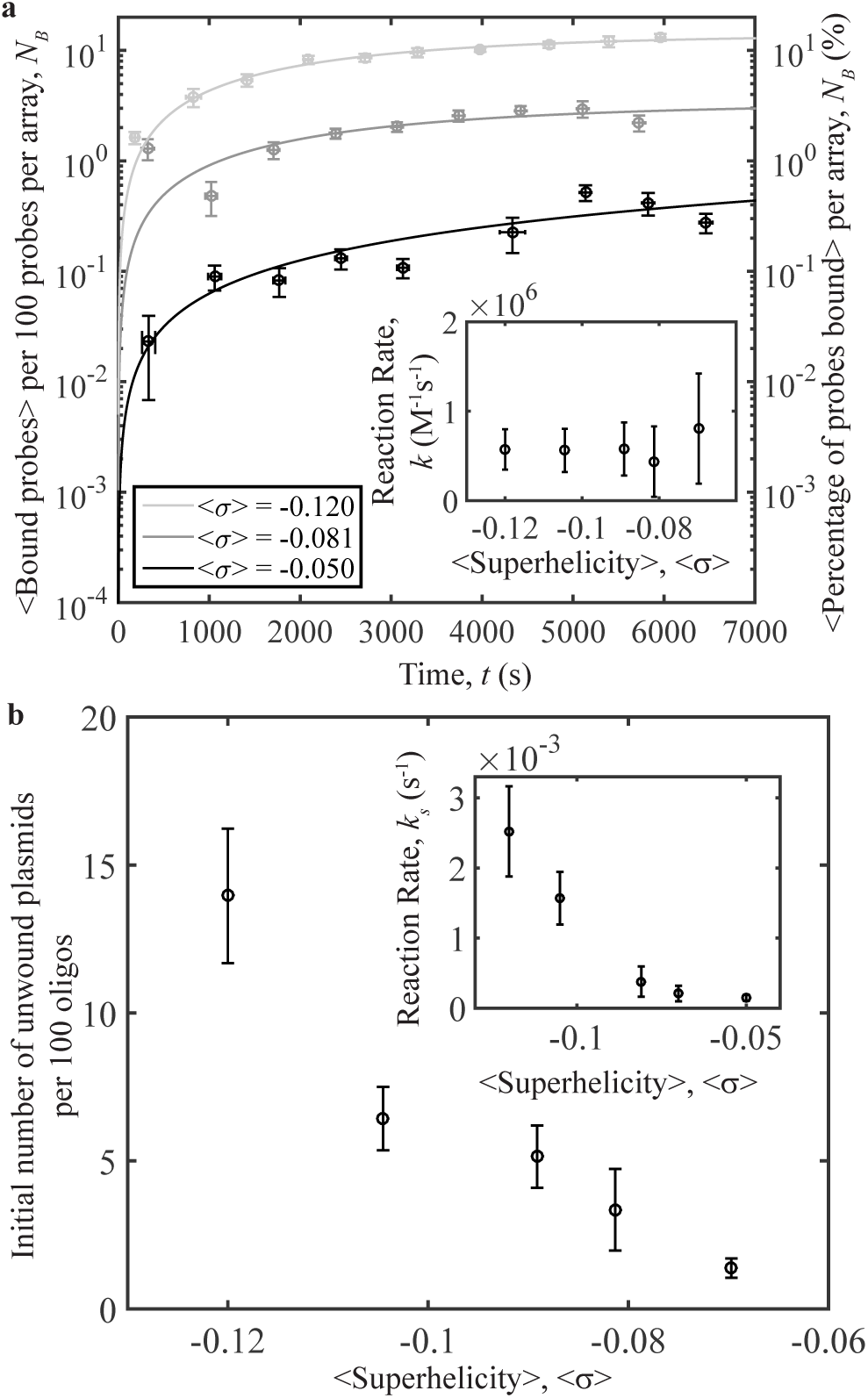
Oligo-plasmid binding and superhelical density. **a)** Number of binding events per 100 probes averaged over 10 different videos vs. time, for a series of < *σ* >. Inset: Reaction rates vs. < *σ* > obtained from the data fits. **b)** The number of unwound plasmids obtained from the fits in a) per 100 probes vs. < *σ* >. Inset: the “short-time reaction rate,” *ks*, vs. < *σ* > for the data in part a. All samples presented here have *sσ* = 0.007. Error bars in a) and the inset of b) denote the standard error of the mean of the binding for the 10 averaged videos on the y-axis, while those on the x-axis represent the standard error of the mean of the time. Error bars on the fit parameters represent the 95% confidence interval of the fit.

### Experimental results: oligo-probe sequence

A series of samples were prepared using oligo-probes of different sizes and/or sequences, and pUC19 with < *σ*>=-0.097 and *s*_*σ*_ = 0.007. Three probes — 20 bases, 30 bases, and 75 bases in length — were designed to bind to the edge of the predicted unwinding region, with 19 bases of each probe chosen to be complementary to the region beside the unwound area. The latter probes with significant predicted overlap with the unwinding region are characterized by similar binding. The shortest probe, with 1-base predicted overlap, has weaker, but non zero binding.

An additional 30-base probe was designed to target the center of the unwinding region (see SI for probe sequences, synthesis, and further discussion). The interactions were similar to those of the 30-base probe targeting the edge (Fig. 7). That the two probes targeting different regions bind nearly as well as one another is interesting: while the unwinding region may be fluctuating in size, there are still enough unwound bases able to nucleate the probe-plasmid binding.

**Figure 7.**
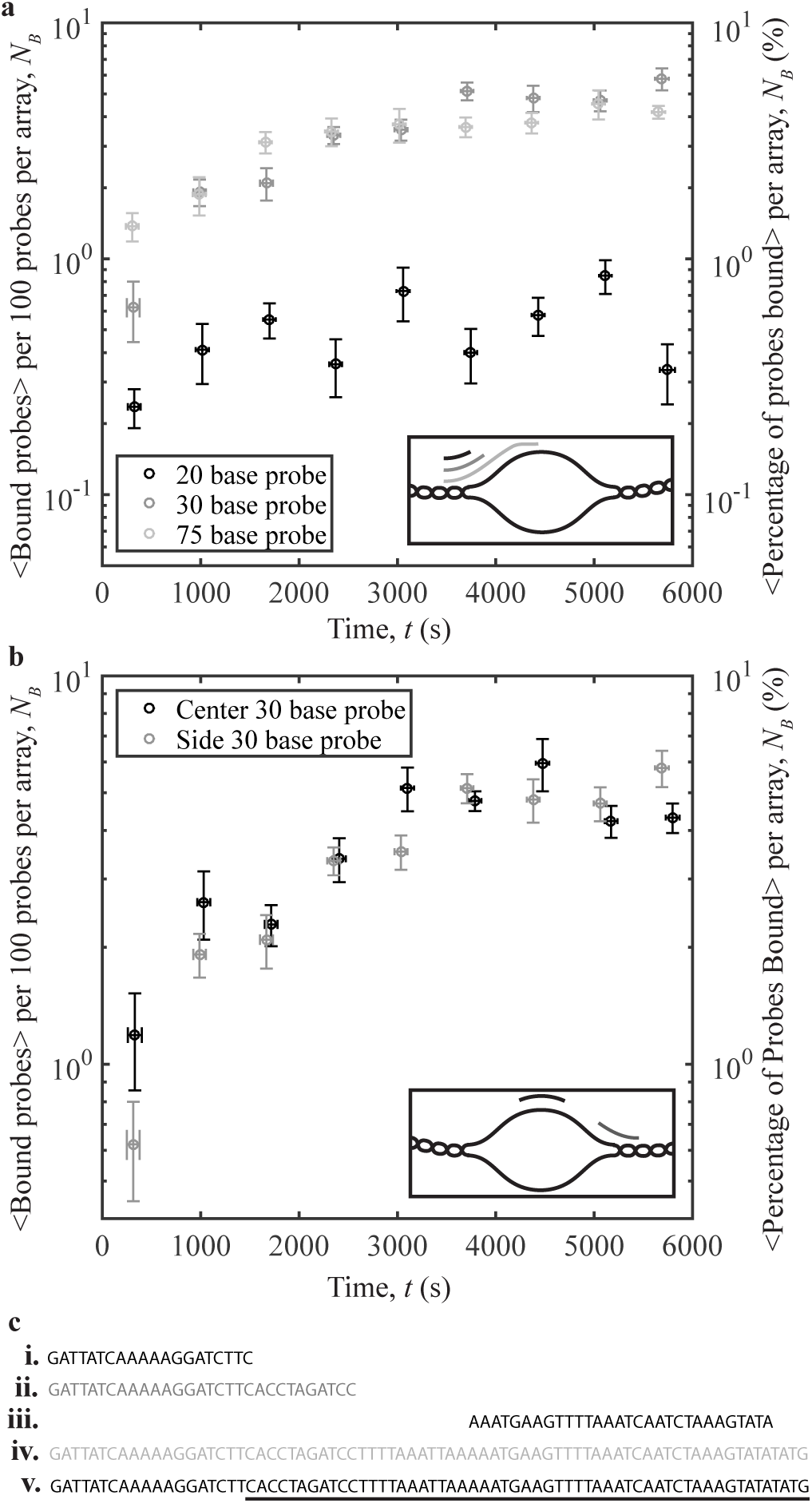
Oligo-plasmid binding and oligo sequence. **a)** Number of binding events per 100 probes averaged over 10 different videos vs. time for 3 different probe sizes: 20 bases (black), 30 bases (dark grey), and 75 bases (light grey). Inset: schematic of the area on the unwinding region where each probe binds. **b)** Number of binding events per 100 probes averaged over 10 different videos vs. time for 2 different probe sequences: probe targeting the center (black), and probe targeting the side (dark grey). Inset: schematic of the area on the unwinding region where each probe binds. **c)** Probe nucleotide sequences for the **i)** 20-base probe, **ii)** 30-base edge probe, **iii)** 30-base center probe, and **iv)** 75-base probe. **v)** shows part of the pUC19 unwinding sequence (underlined), and the wound bases adjacent to this region (not underlined). Error bars on the y-axis denote the standard error of the mean of the binding for the 10 averaged videos, while those on the x-axis represent the standard error of the mean of the time.

These probes are representative of the different possible reactions that occur *in vivo*. Comprehensive knowledge of DNA structure is important for understanding disease. Many cancers are hypothesized to be related to, or caused by, malfunctions in supercoil-induced unwinding (36, 37). In addition, genome editing technologies such as CRISPR-Cas9 can be used to target disease-related regions that participate in strand binding events (38). This technique uses a guide RNA probe that binds to a specific DNA region, conceptually similar to the interaction presented in this work. Our technique may enable advances in CRISPR-Cas9 biotechnology, as well as advance our understanding of transcriptional regulation, DNA replication, and gene editing processes.

### Experimental results: binding kinetics

To gain insight into the timescales of probe/plasmid complex formation, we tracked fluorescence trajectories of single probes for up to 6 minutes and plotted their intensities vs. time. Bound and unbound states were distinguished using the spatial variance in intensity within each pit. For unbound probes, the intensity is uniform across the pit, characterized by a low spatial variance. When bound, the intensity is localized, characterized by a high spatial variance. Though observing capture events are rare, we have successfully found some (Fig. 8). The intensity during such events is characterized by large fluctuations for an extended period, which could reflect transient binding prior to capture, as well as local diffusion by the probe in the vicinity of the unwound strands. Developing and testing mechanistic models of the kinetics of DNA-DNA and protein-DNA interactions, using this methodology, is an exciting research opportunity, which offers important potential contributions towards biomedicine and biotechnology. For more discussion on this method, please refer to SI.

**Figure 8.**
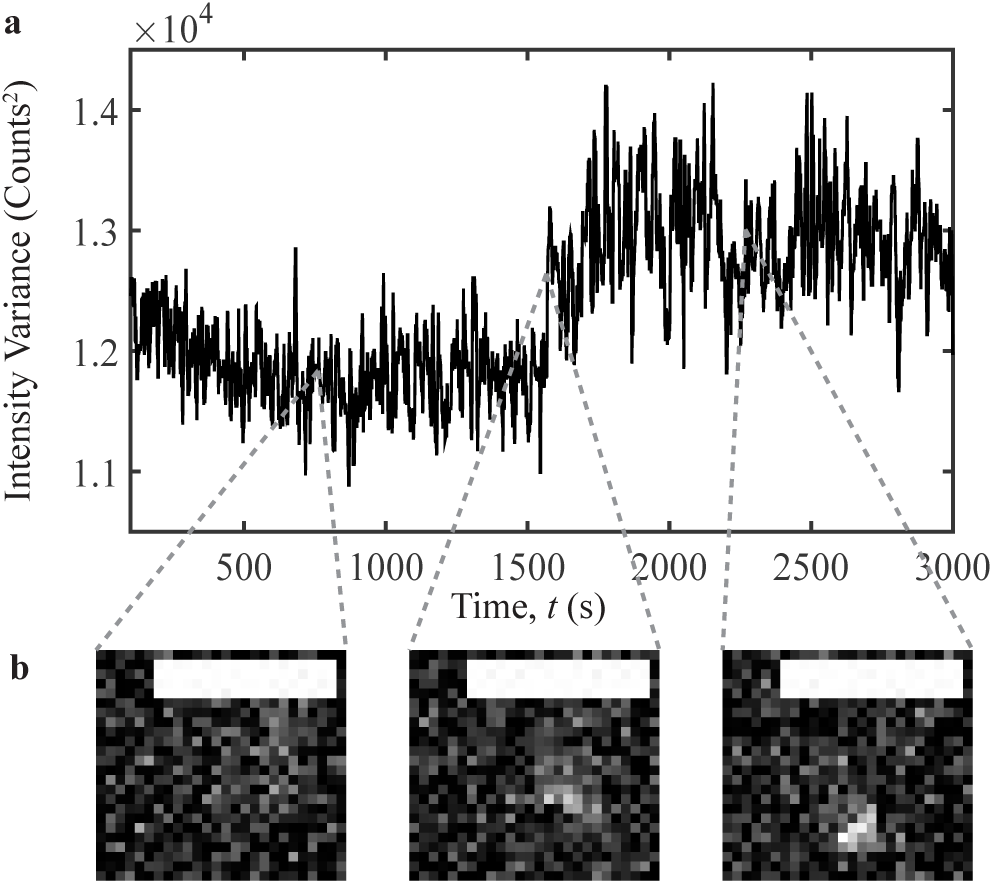
Oligo-plasmid binding capture event. **a)** The spatial variance in intensity vs. time for a probe/plasmid binding event. This event was captured for a plasmid with < *σ* >= -0.151 and *sσ* = 0.007 at T = 31°C. A median filter with a 10-frame window was applied to suppress noise. **b)** Images selected at a series of observation times. Background noise was subtracted using a rolling ball algorithm with a 50 pixel radius. Scale bars denote 2.5 *µ*m.

### Experimental results: topoisomerases IA-driven unbinding

A temperature of ≈60°C is necessary in order for the smallest oligonucleotide probe used to unbind from the unwinding site. As the maximum experimental temperature was 40°C, there was not enough thermal energy to witness significant unbinding of the probes on an experimental timescale. Thus, to demonstrate the reversibility of the reaction, we pre-incubated a sample of supercoiled pUC19 with an average ⟨*σ* ⟩= -0.120 at 37°C until all probes were bound (see SI for further details). After mixing topoisomerase IA with the sample, we inserted it into the microscopy system and counted the number of binding events at different points in time (Fig. 9).

**Figure 9.**
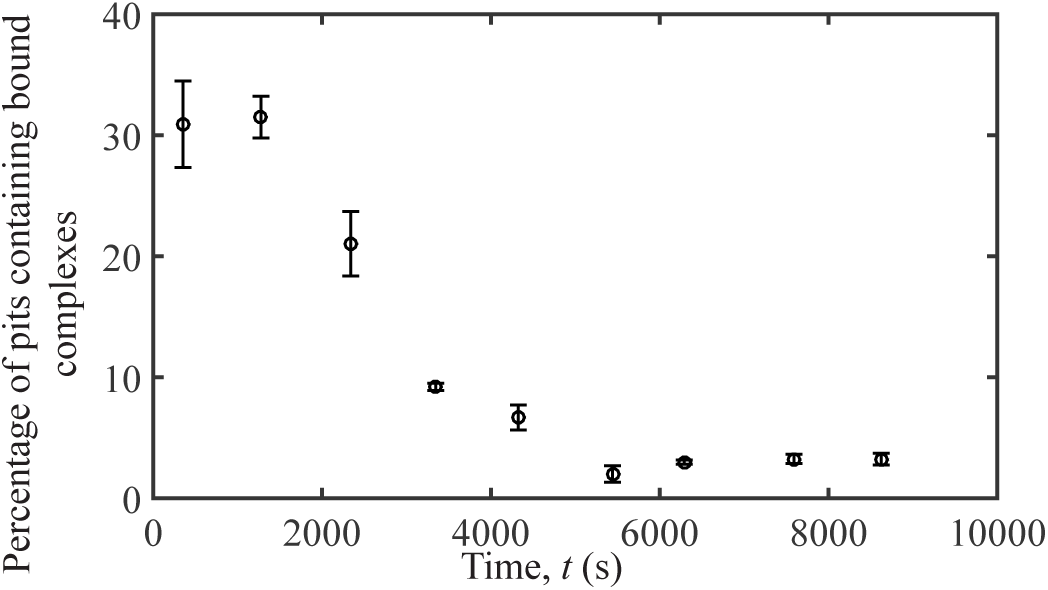
Plasmid-probe unbinding initiated by topoisomerase IA. Percentage of pits containing bound complexes over time for a solution containing 752 pM of the 30-base pair probe and 21.1 nM of pUC19 in 20 mM tris, 50 mM potassium acetate, and 100 *µ*g/mL BSA after the addition of 28.48 *µ*g/mL of *E. coli* topoisomerase IA. Topoisomerase IA was added at *t* = 0.

All probes were bound to plasmids at the start of the experiment. Over time, fewer bound probes and more free probes were observed in each video (Fig. 9). It is interesting to observe that the topoisomerase reaction was slower than expected given that there are, on average, 350 topoisomerase molecules in a single pit.

Unbinding was not observed when supercoiled pUC19 and probe complexes were left on the microscope for comparable periods of time without the addition of topoisomerase IA. This supports that the interaction between topoisomerase IA with the DNA causes the oligo to unbind. The dissociation mechanism may be either through the enzyme interacting directly with the bound probe on the unwinding site and subsequently forcing it off, or the activity of topoisomerase IA itself, namely the relaxing of the supercoiled plasmid, caused the probe to detach from the plasmid, demonstrating reversibility of the reaction.

## DISCUSSION

The experimental results and guiding theory indicate that competing structural transitions play a critical role in supercoil-induced site-unwinding, and subsequent site binding by probe molecules. We find that the interaction is suppressed at lower temperatures, which is consistent with higher predicted instances of Z-DNA formation. This result supports the idea that higher-order DNA structures may play an important role in mediating strand separation, which is of particular interest in the context of DNA replication and repair, and emerging biotechnologies that edit genomes. Specifically, it would apply to CRISPR-Cas9 techniques, as well as providing a better understanding of replication of the plasmids responsible for antibiotic resistance in bacteria.

Our quantitative observations of rare, topology-mediated binding events enable us to establish their strong dependence on temperature, unwinding level, and probe sequence. Simultaneously, we establish a sensitive, open platform capable of visualizing a wide class of weak and transient molecular processes, with single-molecule sensitivity, which depend upon the topological and mechanical properties of molecules. As such, this technique could be applied to study supercoil-induced unwinding in positively supercoiled DNA, and DNA triplexes.

In our experiments, we observe nearly ten times less binding of the “short” (20-base) probe, as compared to the “long” (30-base, 75-base) probes targeting the edge of the predicted unwinding region; however, it is interesting that we observe non-zero binding for the former. In selecting the “short” probe, we sought to measure the degree of binding when introducing only a small (1-base) degree of complementarity between the probe and the edge of the predicted unwinding region. To explain this, consider that while the unwinding region is predicted to be 80 bases long, this is at or above a threshold unwinding probability of five percent; the bases at the edge of the unwinding site have a non zero probability of opening as well. When a 20-base probe binds to the unwinding site, it is likely that one or more of the bases at the edge of the unwinding site are single-stranded and available for binding, allowing for the nucleation and subsequent annealing of the whole probe to the plasmid. In general, our results indicate that even a small predicted unwinding probability (e.g. less than five percent) at the edge of a site can enable the competitive displacement of wound bases on the plasmid, due to the consequent release of mechanical tension within the system and overall reduction in free energy caused by probe/plasmid base-pairing.

Additionally, we observe a similar number of binding events when comparing the 75-base and 30-base probes. The number of binding events for the 30-base probes, which respectively target the center vs. the edge of the predicted unwinding region, do not demonstrate significant deviations in binding when compared to one another. High temperatures and supercoiling levels appear to provide similar binding access to these probe molecules.

Physiological conditions are rarely at equilibrium and inherently complex; thus, single-molecule methods that provide kinetic information and illuminate complex dynamics are important. While equilibrium models have been developed for supercoil-induced unwinding, current methods to predict the kinetics of unwinding are not available for the complex system presented here. Our experimental method provides a direct, empirical means of collecting and statistically assessing dynamic data without mechanically or chemically constraining the molecules, and can inform model development.

Compared to existing alternative methods, confining molecules to pits and imaging with CLiC microscopy is advantageous in that it does not perturb the elastic entropy of the system under observation. By avoiding the chemical or mechanical modifications to the molecules that typically occur in total internal reflection fluorescence (TIRF) microscopy or optical/magnetic tweezers, interactions and reactions can evolve with minimal disturbances introduced by the method itself. Additionally, our method can provide kinetic data on torsionally activated strand separation under a wide range of conditions, particularly non-equilibrium conditions. This is a distinct advantage over studies of supercoiling in gel-constrained DNA, which provide only information on structural states that are visible on a gel: the small quantities of uncommon or rare structures in molecules are unable to be visualized (25).

Our ability to keep freely diffusing molecules under observation for long timescales allows observation of unconstrained structural dynamics and interactions. Although this study focuses on indirectly measuring supercoil-induced DNA unwinding, our technique could be used to investigate its dynamics directly. For example, one could devise a plasmid system in which multiple fluorophores are inserted directly into or near the edges of an unwinding site.

Temporal control over the solution and confinement environment is also possible by extensions of the presented techniques. Further, the capability to fabricate custom pit geometries enables us to optimize the single-molecule imaging conditions for assaying different sizes and concentrations of molecules in solution. Exploring the effects of nanoscale dimensions and molecular crowding is a subject of immense interest as it mimics conditions in the cellular nucleus; these conditions are made accessible by our methods.

Finally, gene editing technologies, such as CRISPR-Cas9, could benefit from a more thorough understanding of nucleotide-nucleotide interactions. The ability of our technique to observe DNA-DNA interactions is quite applicable to those genome editing techniques that make use of a complementary RNA probe to bind to specific DNA regions. Further insight into how these systems function on a molecular level can only help develop these techniques to become more effective.

## CONCLUSION

In conclusion, we have used CLiC microscopy and nanopits to observe and characterize the interactions between a series of fluorescently-labeled DNA probes and a target unwinding region on a supercoiled DNA plasmid. We have shown that by increasing the temperature and level of plasmid supercoiling, we are able to increase the rate of binding of the probes to the target region. By performing experiments as a function of the probe size and sequence, we have discovered that probe molecules are capable of interacting with and binding to the unwinding region even when the probability of base pair denaturation at the target area is low. This result - that even a short probe molecule remains bound - means that the global free energy of the plasmid system is lowered when the unwinding site remains denatured, due to the consequent release of global torsional strain in the DNA molecule. This understanding, consistent with our supporting theory, has implications towards characterizing a wide range of dynamic processes in DNA biophysics, such as DNA replication, where higher order DNA topology plays a critical role; furthermore, it is useful towards developing DNA-based biotechnologies, such as CRISPR-Cas9. What has enabled these contributions is the new method that we present here - the ability to visualize small ensembles of interacting DNA and probe molecules, in large pit arrays, for long times - without tethering them to surfaces or beads. We have shown that we can detect rare kinetic events which are topology-dependent; events which are otherwise out of reach to existing TIRF and tweezer technologies.

## FUNDING

This work was supported by the National Sciences and Engineering Research Council of Canada; the Fonds de recherche du Québec - Nature et technologies; and the Canada Foundation for Innovation.

### ACKNOWLEDGEMENTS

The authors thank the Canadian Foundation for Innovation, the National Science and Engineering Research Council of Canada Discovery program, The Fonds de recherche du Quebec Nature et technologies (FRQNT) Team Grant and Fellowship programs, the Cellular Dynamics of Macromolecular Complexes CREATE fellowship program, the Bionanomachines fellowship program, and McGill University for funding and resources. C.S. thanks CMC Microsystems for fellowship funding for fabrication. The authors thank Gil Henkin and Frank Stabile for fabrication support; Susi Kaitna for training in plasmid transformation; Alec Silver for early sample preparation; Chris Cayen-Cyr for early analysis work; and Tom Edwardson and Hanadi Sleiman for oligo purification on an HPLC. Further, Stephen Michnick contributed important manuscript feedback and scientific discussions.

### Conflict of interest statement

None declared.

