## Supplementary Materials for "Visualizing structure-mediated interactions in supercoiled DNA molecules"

### Supplementary Information (SI): Visualizing structure-mediated interactions in supercoiled DNA molecules

#### 1 pUC19 Purification and Supercoiling

##### 1.1 Plasmid Purification

Samples of pUC19 (catalog # N3041L) were ordered from New England Biolabs (Whitby, ON). A diluted solution was transformed into competent DH5$\alpha$ cells (catalog # 18265-017) from Life Technologies (Waltham MA) and grown on an agar plate overnight at 37°C. One colony was chosen from the plate and grown in Luria broth (LB) at 37°C in a shaking incubator until an absorbance value at 600 nm (or optical density, OD) of 0.4 was obtained. 700 $\mu$L of cells in media were diluted with 800 $\mu$L of 40% glycerol, then flash frozen and stored in a -80°C freezer. Subsequent stocks of cells were grown from this stock in LB media at 37°C overnight and purified using a Geneaid Plasmid Midi Kit (catalog # PI025) from Frogga Bio (North York, ON).

Plasmid samples suspended in 10 mM tris (pH 8.0) were quantified by spectrophotometry, aliquoted, and stored in a freezer at -20°C. Plasmid sample purity was assessed by DNA electrophoresis using gels made of 1% agarose, 10 mM tris base (pH 8.0), 20 mM acetic acid, and 1 mM EDTA in an Owl EasyCast B2 from ThermoFisher Scientific. Gels were run at room temperature for 1 h at 120 V, stained with SYBR Safe (catalog # S33102) from Life Technologies (Waltham, MA), and imaged using a GelDoc EZ Imaging System (catalog # 1708270) from Bio-Rad (Mississauga, ON).

##### 1.2 Topoisomer Production

Plasmid samples were supercoiled following the procedure by Keller [1] wherein increasing amounts of ethidium bromide (EtBr) are mixed with a fixed amount of pUC19 and topoisomerase IB to create solutions of topoisomers. Calf thymus DNA topoisomerase IB (catalog # 38042024) was purchased from Thermo Fisher Scientific (Toronto, ON). The reaction was carried out in 50 mM tris-HCl (pH 7.5), 50 mM KCl, 10 mM $MgCl_2$, 0.1 mM EDTA, 0.5 mM DTT, and 30 $\mu$g/mL BSA for 2 h at 37°C. As the amounts of EtBr added to plasmid solution produced positive supercoils, it was necessary to use topoisomerase IB, which can relax both positive and negative supercoils, and not topoisomerase IA which relaxes only negative supercoils. All reactions contained 10 $\mu$g of pUC19 and 10 units of topoisomerase IB in a total volume of 100 $\mu$L and were stopped by heating the samples for 20 min at 65°C.

The DNA was precipitated via ethanol precipitation in order to remove salts. Contaminant proteins were degraded by reacting the sample for 30 min at 25°C with Proteinase K (catalog #P8107S) from New England Biolabs (Whitby, ON) in the presence of 0.5% SDS. The samples were further purified following this reaction using a Qiaquick Gel Extraction Kit (catalog # 28704) from Qiagen Inc. - Canada (Toronto, ON), suspended in 10 mM tris, and stored in a freezer at -20°C.

Topoisomer superhelicities were checked by gel electrophoresis using 2% agarose gels with 10 mM tris base (pH 8.0), 20 mM acetic acid, and 1 mM EDTA in an A5 Owl EasyCast gel system (catalog # 27372-134) from VWR International (Ville Mont-Royal, QC). To visualize the different distributions of topoisomers, both the gel and the buffer were supplemented with 3 mg/L chloroquine diphosphate, 6 mg/L chloroquine diphosphate, or 6 mg/L chloroquine diphosphate and 0.1 mg/L EtBr (SI Fig. 1). Electrophoresis took place at room temperature for 40 h at 30 V. Gels were stained in a 1% (v/v) SYBR Safe bath for 30 min and imaged using a GelDoc EZ Imaging System (catalog # 1708270) from Bio-Rad (Mississauga, ON).

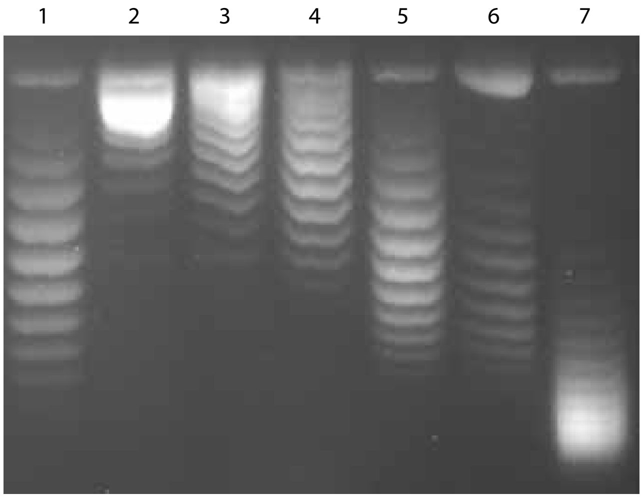


**Supplementary Figure 1: Topoisomer separation using gel electrophoresis.** Gel electrophoresis of topoisomer samples with 6 mg/L chloroquine diphosphate and 0.1 mg/L EtBr. Each lane contains a mixture of topoisomers centered at a $\sigma$ of –0.152 (lane 1), –0.133 (lane 2), –0.121 (lane 3), –0.109 (lane 4), –0.098 (lane 5), –0.082 (lane 6), and –0.07 (lane 7).

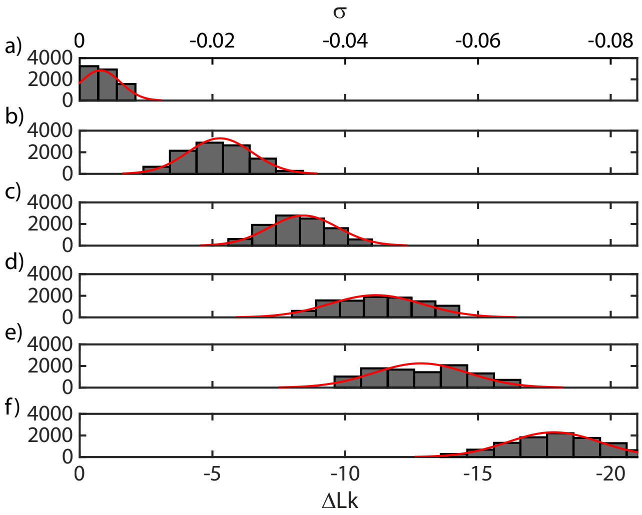


**Supplementary Figure 2: Mean topoisomer determination.** Gaussian distribution of topoisomer samples centered at estimated superhelicities of **a)** $\langle\sigma\rangle = 0.000$, **b)** $\langle\sigma\rangle = -0.020$, **c)** $\langle\sigma\rangle = -0.031$, **d)** $\langle\sigma\rangle = -0.043$, **e)** $\langle\sigma\rangle = -0.055$, and **f)** $\langle\sigma\rangle = -0.07$.

#### 1.3 Topoisomer Gaussian Distribution

Topoisomer samples produced from cells naturally contain a Gaussian distribution of superhelicities, as demonstrated in our samples. The distribution in each sample was quantified using ImageLab software (Bio-Rad), as in SI Fig. 2. Each topoisomer band was manually selected to exclude nicked DNA and dimers, which were also present. The band intensities were fitted to a Gaussian distribution using Matlab. With the calculated distribution, we were able to quantitatively compare the theory to experiments; for example, the plots of predicted unwinding as a function of $< \sigma >$ take into account each sample's inherent Gaussian superhelicity distribution $s_\sigma$. The determined fit values for the mean superhelicity $< \sigma >$ and standard deviation $s_\sigma$ were used to create the plots in this work.

### 2 Oligonucleotide Probe Synthesis and Labeling

Using the algorithm published by Zhabinskaya *et al* [2], Site 1 and the bases before and after it were determined to be:

5' - GAG ATT ATC AAA AAG GAT CTT CAC CTA GAT CCT TTT AAA TTA AAA ATG AAG TTT TAA ATC AAT CTA AAG TAT ATA TGA GTA AAC TTG GTC TGA CAG TTA CCA ATG CTT AAT- 3'

where the underlined sequence denotes the predicted unwound bases at 37°C, 22.5 mM ionic strength, and $\sigma = -0.055$.

Unlabeled single-stranded oligonucleotide probes were purchased from Integrated DNA Technologies (Coralville, IA). An amine modification was added to the 5' end to allow for fluorophore labeling using Cy3B dyes conjugated with an NHS ester. The DNA sequences ordered are given below.

**20-base probe:** 5' - /5AmMC6/GA TTA TCA AAA AGG ATC TTC - 3'

**30-base probe:** 5' - /5AmMC6/GA TTA TCA AAA AGG ATC TTC ACC TAG ATC C - 3'

**30-base center probe:** 5' - /5AmMC6/AA ATG AAG TTT TAA ATC AAT CTA AAG TAT A - 3'

**75-base probe:** 5' - /5AmMC6/GA TTA TCA AAA AGG ATC TTC ACC TAG ATC CTT TTA AAT TAA AAA TGA AGT TTT AAA TCA ATC TAA AGT ATA TAT G - 3'

The first three probes are complementary to one end of the dominant unwinding region (Site 1) on pUC19, while the 75-base probe is complementary to the predicted unwinding region's end and center. Only one probe was designed to exclusively target the center, as our numerical simulations predict that unwinding nucleates at its center. The only limitation on binding in this region therefore rests entirely on whether the site is unwound or not, meaning the size of the oligo probe will not make a difference as they will all target this same unwound region.

Extra bases corresponding to the region just beyond the predicted unwinding site were added to aid in binding.

Cy3B NHS ester (catalog # PA63101) was purchased from GE Healthcare (Mississauga, ON). The Cy3B NHS ester was dissolved in dimethyl sulfoxide (DMSO) at a concentration of 1 mg/mL and reacted with the DNA overnight on a rotating platform. The resultant reaction was alcohol precipitated using ammonium acetate and ethanol a total of 3 times to remove unreacted dye. To purify labeled probes from unlabeled probes, the DNA sample was run on a C18 reverse phase column using high performance liquid chromatography (HPLC).

### 3 Flow Cell Preparation

Coverslips were cleaned using the procedures described in Berard *et al* [3]. The flow cells were assembled by taping together the top (No. 1.5, with pits) and bottom (No. 1, with inlet and outlet holes) surfaces, using 10-$\mu$m- or 30-$\mu$m-thick laser-cut double-sided tape gaskets, as described in the same reference. The bottom surface of the flow cells contained square arrays of 27,556 3-$\mu$m-diameter pits (1 mm $\times$ 1 mm), which were photolithographically defined on No. 1.5 D263 glass coverslips (catalog # CA48366-249-1) purchased from VWR (Radnor, PA) and etched to a depth of 500 nm by reactive ion etching. The top surface of the flow cells contained inlet and outlet holes, which were drilled in No. 1 coverslips (catalog # CA48366-089-1) from VWR International (Ville Mont-Royal, QC).

### 4 Convex Lens-induced Confinement (CLiC) Microscopy

The Convex Lens-induced Confinement (CLiC) device described in Figure 1 of Berard *et al* [3] was used for loading and imaging samples. First, the sample was injected into a custom microfluidic chuck, which loaded it into a glass flow cell. After the sample was loaded into the flow cell, the CLiC device mechanically deformed the roof of the flow cell, lowering the top surface into contact with the bottom surface, which contained the embedded microscale pits. Molecules were thereby loaded into these pits for observation, from the top. The pits were sealed, as verified by microscopy, since the glass surrounding the pits was pressed into mechanical contact. Importantly, molecules as small as single oligos and single fluorophores do not escape the pits when the surfaces are pressed together; their imaged fluorescence remains confined within the pits.

### 5 Temperature Control

A temperature control system that uses two heaters was developed for this work, with one heater integrated directly behind the CLiC lens, and a second heater wrapped around the objective directly below the sample, as in SI Fig. 3. This configuration minimizes temperature gradients across the sample, providing the temperature accuracy required for the presented studies.

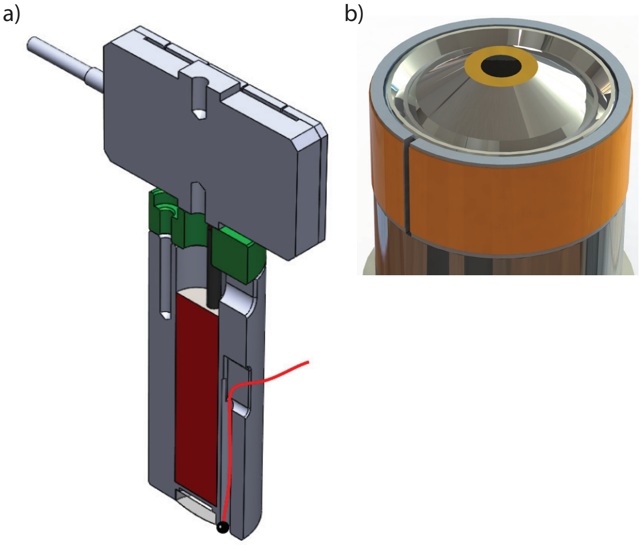


**Supplementary Figure 3: Schematic of two-part implementation of temperature control. a)** Lens heater and thermistor. The aluminum tube on which the CLiC lens is mounted contains a cartridge heater (red cylinder). A thermistor (red wire and black bead) located beside the CLiC lens senses the temperature at the lens position. A spacer thermally insulates the aluminum tube from the piezoelectric actuator (top grey rectangular component). **b)** Objective heater. Adhesive-backed kapton strip heaters mounted to a thin aluminum split-ring collar are used to heat the objective. A thermistor mounted within the collar provides temperature feedback.

Small temperature changes can cause nanoscale thermal expansion of the CLiC device components and microscope objective, potentially varying the confinement within the flow cell. To ensure stable temperature control, we developed a custom proportional-integral-derivative (PID) temperature controller. Temperatures at both the CLiC lens and objective were measured using thermistors (Amphenol MC65F103, $\pm 0.1°$C tolerance) located beside the lens and within the objective heating collar. Temperature fluctuations with this system typically did not exceed $\sim \pm 0.05°$C.

The possible temperatures for our system ranged between 28°C - 40°C. Ambient temperatures for the microscope ranged between 26°C - 28°C, meaning the minimal stable temperature at which the device could operate was 28°C. Regarding the maximum temperature available, the Nikon objective temperature is only rated to operate up to a temperature of 37°C. The reason for this is two-fold. First, the adjustable temperature collar on the objective can only correct for temperatures up to 37°C, meaning image quality deteriorates with increased temperature above this level rendering images above this out of focus. Second, the epoxy in the objective that holds the optical components together melts at temperatures above 40°C, meaning that heating above this level will damage the equipment.

### 6 Microscopy Procedure

Prior to sample injection, NF oil (catalog # MXA22024) from Thorlabs, Inc. (Newton, NJ) was placed on the 100× objective. The custom temperature controller was set to the desired temperature for the experiment. A glass coverslip was placed on an aluminum sample plate and the objective was brought to the focal height used during a CLiC experiment. Using a custom-made dichroic cube (532/647) from Chroma Technology Corp. (Bellows Falls, VT) that reflects 532 nm laser light and a power meter (catalog # PM100D) from Thorlabs, Inc. (Newton, NJ), the transmitted laser power through the coverslip was measured to be ˜30 mW. After measuring laser power, the flow cell and chuck were installed.

Prior to sample insertion, 5 $\mu$L of a buffer solution of 12 mM tris, 25 mM HEPES, and 10 mM NaCl at pH 8.0 was inserted into the flow cell to wet the chamber. Using this buffer, the height at which the CLiC lens presses the top and bottom coverslips into contact was determined. The x-y position of the contact point for the CLiC lens was also determined. If the contact point for the two coverslips was not directly on the pit array, the lens was lifted by 20 $\mu$m, the flow cell position under the CLiC lens was adjusted using the device's XY translation stage. This stage moved the flow cell with respect to the CLiC lens and objective, allowing control over the contact point between the lens and the flow cell. Once these positions were co-aligned, the majority of the buffer was removed using a pipette.

Samples were prepared in a solution characterized by 12 mM tris, 25 mM HEPES, 10 mM NaCl, 2.5 mM protocatechuic acid (PCA), 50 nM protocatechuate 3,4-dioxygenase (PCD), and pH of 8.0. This corresponds to an approximate overalll ionic strength of 22.5 nM. The pUC19 concentration was 21.1 nM and probe concentration was 752 pM. Once confined in the chamber, the sample was effectively isolated from the outside world over the time period of the observations. This is consistent with minimal photobleaching of the fluorophores being observed, due to both the effectiveness of the "oxygen scavengers"' (PCA, PCD) at depleting the initial oxygen from the solution, as well as the suppression of oxygen exchange through the non-porous glass walls.

For each sample, 90 videos were acquired at ≈1 min intervals with 50 frames at 50 ms exposure/frame using an Xion Ultra 888 camera (Andor) and a CFI Apo TIRF 100× objective from Nikon Canada (Mississauga, ON). A series of images was acquired with 2040 frames at 50 ms exposure/frame using this setup, to form a video. Between each video, the CLiC-lens was lifted by approximately 10-20 $\mu$m and oscillated slightly to allow the solution in the pits to be exchanged with the molecules diffusing freely in solution, before the system was resealed. In this way, each video represents an experiment performed on new molecules. Importantly, the ability to replenish the pits with fresh sample and acquire new data enabled high statistics to be obtained.

### 7 Experimental Verification of Plasmid Concentration

The average number of pUC19 plasmids per pit was verified using the experimental microscopy procedure (described above) and pUC19 samples labeled with YOYO-1 (catalog # Y3601, Fisher Scientific, Toronto, ON) at approximately one fluorophore per ten base pairs. The YOYO-1-stained plasmids were used for establishing per-pit plasmid number so that the per-plasmid fluorescence signal would be large and insusceptible to photo bleaching. 70 $\mu$L of 1.055 nM YOYO labeled pUC19 in 1×TE buffer and 1% BME (catalog # M3148-25ML, Sigma-Aldrich, Oakville, ON) was inserted into a flow cell and trapped in pits using the CLiC device. The sample was illuminated with a 488 nm laser at 38.59 $\mu$W passing through a custom 488/647 dichroic cube from Chroma Technology (Rockingham, VT). Five 200-frame videos were recorded at 100× magnification, and the sample was replenished between videos by lifting and oscillating the CLiC lens. Single plasmids were visible as diffusing 'particles' in each pit and the number of plasmids per pit was counted. The mean number of plasmids per pit was found by fitting a Poisson distribution to a histogram of the count. For the sample dilution used in this counting experiment, there were 1.19 ± 0.03 plasmids per pit. Extrapolating from this value, there were 23.8 ± 0.6 plasmids per pit at the experimental concentration of 21.1 nM.

### 8 Experimental Microscopy Controls

It is possible that the probe could forcibly anneal to a region on the pUC19 plasmid for which it has only partial complementarity. To control for this, we tested probe binding of the 75-base probe with a control plasmid, pMAL-pIII (catalog #N8101S) from New England Biolabs (Whitby, ON). Low complementarity between the plasmid and probe indicate that little binding, if any, should occur. Following the same microscopy procedure as above, we found only 3 binding events over an experimental time frame of ˜2 h (data not shown). This supports that the binding that we are observing in our studies is predominantly occuring in the intended region complementary to the probe sequence.

To further control for other possible forms of binding, we tested a probe design from the literature [4] (5' - /Cy3/ - AGA CTA GAC CCC AAA AAA AAA AAA AAA AAA -3'), and ordered it from IDT. This probe has little complementarity to Site 1 on pUC19, but is similar in size to the probes used during our experiments. Using the microscopy procedure described above with 21.1 nM pUC19 at $\sigma = -0.055$ and 752 pM of this probe, we observed no binding events over all movies. This further supports that the binding that we observe in our studies is due to complementary annealing of our probes to Site 1 on pUC19.

High laser powers, such as those used in these experiments, could be conjectured to cause photonicking in supercoiled DNA, potentially relaxing the supercoiling and preventing unwinding. Were this to be the case, the probability of photonicking would increase with laser illumination time and would result in reduced observed binding. To control for potential photonicking, a sample of 21.1 nM pUC19 with $\sigma = -0.055$ and 752 pM of long probe was placed inside the CLiC device. The temperature was then increased stepwise, without illuminating the molecules with the laser. Once 40°C was achieved, the sample was left to incubate for a time equivalent to an experiment. 20 videos were then acquired following the experimental microscopy procedure. The amount of binding in this control was found to be slightly less than when the laser power was on, indicating that photonicking is not an issue: if photonicking were influencing results, we would expect to observe more binding complexes with the laser off.

To test our assumption that probe binding can saturate in our system, a sample of 21.1 nM pUC19 at $\sigma = -0.081$ mixed with 752 pM of long probe was incubated overnight at 37°C. After 18 h, the sample was observed at 37°C by following the experimental procedure. All observed probes were bound to plasmids. The temperature was lowered to 27°C and the sample was allowed to incubate on the microscope for 0.5 h. After incubation, all observed probes were still bound. Given sufficient time, we observe the binding to saturate at the experimental temperatures used in this study.

Finally, to verify that the observed binding of the 30-base probe is indeed to the unwinding site on pUC19 and not to another region, we performed an experiment as per the standard microscopy procedure described in SI section 6 using a plasmid with $\sigma = 0$. No binding events were observed over the course of the experiment, indicating that the site must be supercoiled in order to unwind and allow for binding to occur.

### 9 Experimental Verification of DNA Secondary Structures

To verify the presence and locations of secondary structures in pUC19, we used a potassium permanganate fingerprinting assay. Potassium permanganate oxidizes single-stranded thymine bases in DNA, thus preventing them from reannealing. Since DNA secondary structures have single-stranded regions at the transition from B-DNA, potassium permanganate may be used to map out where the secondary structures occur along a given DNA molecule.

To test pUC19 molecules for the presence of secondary structures, pCu19 with a range of superhelical densities were treated with 7 $\mu$L of 100 mM $KMnO_4$ in 20 mM tris and 10 mM NaCl and incubated for 2 min at either room temperature or 37°C. The $KMnO_4$ reaction was quenched with 10 $\mu$L of 14 M $\beta$-mercaptoethanol (BME). Single-stranded regions of the plasmid were cut using S1 nuclease and subsequently cut again with either PciI or HindIII. Both these restriction endonucleases target a single, exact position in pUC19, allowing the positions of non-B-DNA secondary structures to be determined. The samples were run on an agarose gel to measure the size of the resulting DNA fragments and map out alternate DNA secondary structure locations (Fig. 4).

For this protocol, if the two known unwinding sites were the only secondary structures present in pUC19, we would expect to see bands at 1860 bp and 740 bp for the PciI-cut pUC19, and bands at 1500 bp and 1100 bp for HindIII-cut pUC19. These bands correspond to the fragments left after the unwinding site was cut. An additional band at 2686 base pairs, belonging to pUC19 with no secondary structures, should be visible as well. At minor supercoiling levels and room temperature, only linearized pUC19 bands were present, suggesting no secondary structures in the plasmid at these conditions (Fig. 4a and b). For samples with more supercoiling ($\sigma < -0.055$) at room temperature, a number of new bands appeared, indicating the presence of DNA secondary structures. While some bands corresponded to the known unwinding sites, the appearance of a number of other bands indicates other regions with DNA secondary structures.

As unwinding is more likely at higher temperatures than Z-DNA, gel bands caused by Z-DNA that were apparent at room temperature would be gone for the gel with the reaction run at 37°C. Observing the gel of the $KMnO_4$ reactions at 37°C, it is clear there is a reduction in the number of bands compared to the gel of the

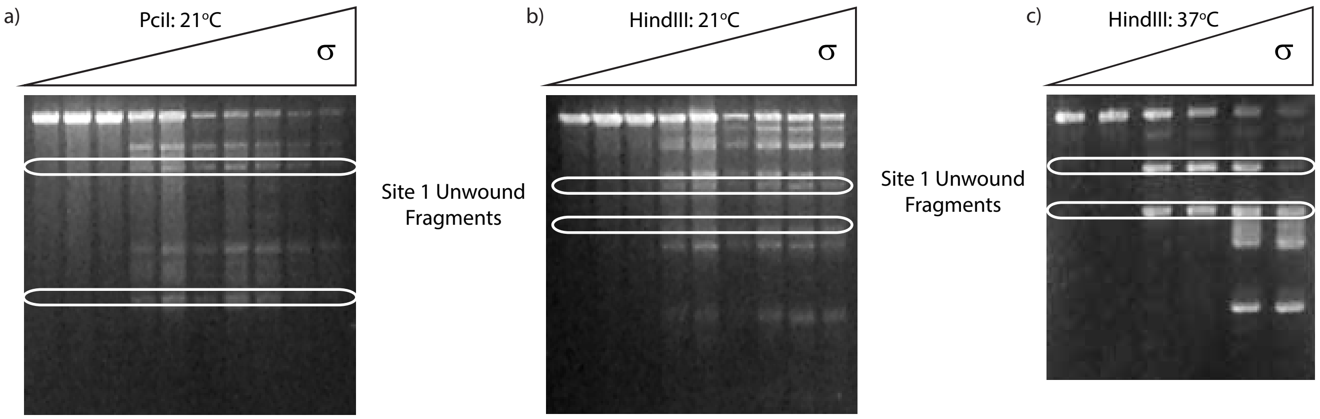


**Supplementary Figure 4: Gels demonstrating DNA secondary structure using $KMnO_4$ fingerprinting.** pUC19 with a supercoiling range of $0 \geq \sigma \geq -0.147$ reacted at room temperature with $KMnO_4$, then with either **a)** PciI, or **b)** HindIII. Following this, they were all treated with S1 nuclease. The white circle indicates the bands corresponding to the Site 1 unwinding site, while all other bands either correspond to pUC19 with no other DNA secondary structures (top band of the gel), or some other structure such as Z-DNA. **c)** Supercoiled pUC19 reacted with $KMnO_4$ at 37°C, then cut with HindIII and S1 nuclease. The pairs of bands appearing here are those of the Site 1 unwinding site, circled in white, and those of pUC19's secondary unwinding site associated with its ampicillin resistance gene. Comparing c) with a) and b), it is evident there are less bands at higher temperatures, indicating the dominance of unwinding at these conditions.

same reaction performed at room temperature (compare Fig. 4a and b with c). The sole exceptions to this are the two pairs of bands corresponding to the two known unwinding regions, one pair corresponding to the unwinding site studied in this work appearing at lower superhelical tension, the second only appearing after sufficient supercoiling ($\sigma \leq -0.074$). These tests provide evidence of Z-DNA at lower temperatures that disappear at higher temperatures in favor of DNA unwinding, supporting the trends we observe in the microscopy data presented in this work.

### 10 Fluorescence Image Analysis Methods

Custom image processing methods and algorithms were established to identify binding events. A supervised Machine Learning (ML) classifier was used to aid in identification of binding events. The analysis steps are described here.

#### 10.1 Locating the Pits

The first step of the image processing procedure identified the pits in which the molecules were trapped so that each one could be checked individually for binding events. For each video, the intensity values for every frame were averaged to get a collapsed image. The background illumination was calculated by performing a convolution of the image with a normalized uniform square matrix, and was subsequently subtracted from the image. Each video thus had a corresponding collapsed image on which the following analysis could be performed.

The positions and spacings of the pits were determined by applying a series of cross-correlation methods with manual user input. A single pit, two adjacent horizontal pits, and two adjacent vertical pits were cropped using a custom graphical user interface (GUI). Next, the single pit was cross-correlated with each set of adjacent pits. The distance between adjacent peaks in these cross-correlations was taken as the spacings between pits in the horizontal and vertical directions, which were then used to estimate the number of rows and columns in the array of pits.

A second cross-correlation was performed between the single pit and the entire collapsed image. The resulting peaks approximately corresponded to the location of individual pits. To fill in missing pits and improve accuracy in positions, a radon transform was performed which identified straight lines connecting the peaks (SI Fig. 5). The intersections of the lines were taken to be the pit positions. Finally, to reduce error due to edge effects, the pits in the left-most and right-most columns, as well as the top and bottom rows, were removed manually.

#### 10.2 Using a Diffusion Coefficient Estimator

As the probe molecules were confined to pits of finite area, their mean squared displacement curves saturated as a function of time. At short times, their mean squared

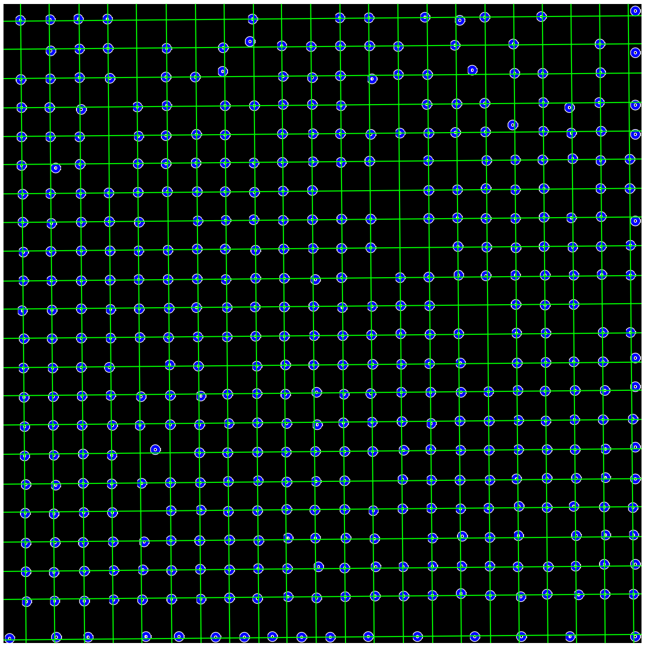

**Supplementary Figure 5: Radon transform for pit localization.** Visualization of the lines (white) detected by the radon transform. The cross-correlation peaks are shown as blue circles.

displacement increased approximately linearly with time. For the purposes of this analysis, it was sufficient to calculate an estimator for the diffusion coefficient, $D_{est}$, using the first point of the plot of mean squared displacement versus time. This is because the large probe-plasmid complexes diffused sufficiently more slowly than the small probes, to distinguish between the two states of the probes. (We note that the reported values of $D_{est}$ are not the true diffusion coefficients, due to the bias introduced by using only the first point of the MSD curve). We begin by addressing the diffusion coefficient estimator in the context of the bound probe-plasmid complexes. To calculate $D_{est}$, the estimate for the diffusion coefficient, the precise location of the probe, which appeared as an area of bright pixels, was tracked for each frame of the video. First, an approximate location of the bright area was found iteratively, using a 2-dimensional sliding window of 6 pixels by 6 pixels. The brightest window within the pit was assumed to be the general location of the fluorescent probe. More formally, the top left corner of this window, $(i_{max}, j_{max})$, was found by:

$$(i_{max}, j_{max}) = \underset{1 \le i \le m-5, 1 \le j \le n-5}{\operatorname{argmax}} \left( \sum_{i}^{i+5} \sum_{j}^{j+5} I(i,j) \right) \quad (1)$$

where $I_{mn}$ was a matrix representing all pixel intensities in the pit.

A flood-fill algorithm was utilized to find the precise outline and center of the bright area. First, the brightest pixel within the defined window was selected as the seed pixel. Next, the component was expanded from the seed in a four-neighbor recursive manner using a traversal technique as outlined in the pseudocode below. The termination threshold was determined empirically as the $85^{th}$ percentile of the $I_{min}$ pixel intensities. The resulting connected component we interpreted as the shape of the probe-plasmid complex, which allowed the calculation of a centroid location. The flood fill algorithm was defined as the following:

**procedure** FLOOD-FILL(seed $[i, j]$, threshold $I_{min}$)
    **if** $[i, j]$ is outside of pit border **then return**
    **else if** $I[i, j] < I_{min}$ **then return**
    Explore [i,j]
    FLOOD-FILL( [i-1,j] , $I_{min}$)
    FLOOD-FILL( [i+1,j] , $I_{min}$)
    FLOOD-FILL( [i,j-1] , $I_{min}$)
    FLOOD-FILL( [i,j+1] , $I_{min}$)

Refer to SI Fig. 6 for the output of this algorithm.

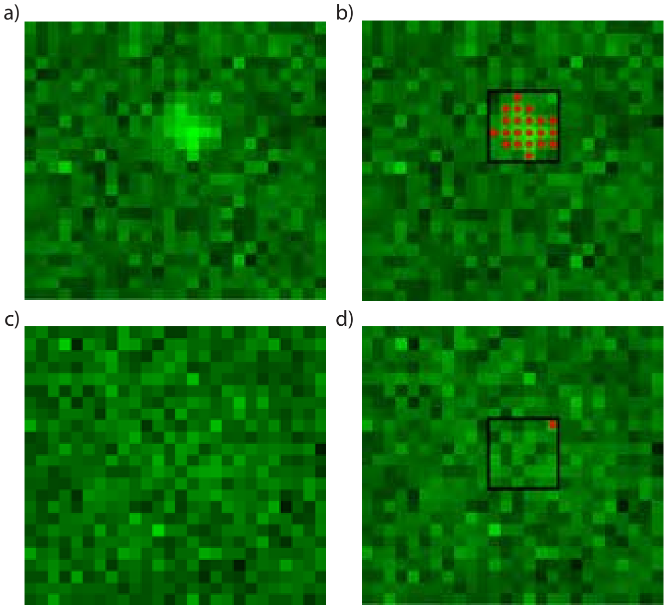


**Supplementary Figure 6: Probe-plasmid complex detection. a)** Raw, single-frame image of a probe-plasmid complex. **b)** A bound probe detected by our algorithm. Red dots illustrate the connected components found by the flood-fill algorithm. The black square illustrates the sliding window. **c)** Raw, single-frame image of an unbound, freely diffusing probe. **d)** As intended for an unbound probe, the flood-fill algorithm does not find any connected components, as demonstrated by the lack of red dots.

The displacements of the particle across two consecutive frames were obtained from the centroid locations of the probe in each frame. Let $S_i$ be the displacement in the $x$ direction from frame $i-1$ to $i$, then the variance of the distribution of step sizes obtained from a video with $K$ frames is:

$$\langle \Delta S^2 \rangle = \frac{1}{K} \sum_{i=2}^{K} (S_i - \mu)^2$$

.

If an unconfined particle takes $N$ steps, then the mean squared displacement is:

$$\langle \Delta X^2 \rangle = N \langle \Delta S^2 \rangle = 2DN\Delta t = 2Dt$$

.

Since confinement within the pit affects the displacements when multiple steps were taken, we use the case where $N = 1$ (equivalently $t = 1$). In this case, $\langle \Delta X^2 \rangle = \langle \Delta S^2 \rangle$. This was repeated for the y direction; the estimated two-dimensional diffusion coefficient was obtained by:

$$D_{est} = \frac{1}{4} \langle \Delta r^2 \rangle = \frac{1}{4} (\langle \Delta S_X^2 \rangle + \langle \Delta S_Y^2 \rangle)$$

Next, we address the diffusion coefficient estimator in the context of the unbound probes. The unbound probes were able to diffuse everywhere inside the pits within the 50 ms exposure time; thus there were no "concentrated bright spots" inside the pit to track using the above algorithm. Rather, the above code detected random bright sections of the diffuse intensity spread across the pit. The random selection of bright points produced an estimated diffusion coefficient much higher than that for the bound probes (SI Fig. 7a). In this way, the above methodology was sufficient to distinguish between the unbound probes and the bound probe-plasmid complexes.

#### 10.3 Calculating the Variance in Location

Occasionally, fluorescent probes would stick to the glass coverslips. In order to differentiate stuck molecules from diffusing molecules, such as the probe-plasmid complexes of interest, the variance in the explored locations, $V$, was calculated. This informally corresponds to the "randomness" of the movement of the probes. Across $k$ frames:

$$V = \sum_{t=1}^{50} \frac{(x_t - \bar{x})^2 + (y_t - \bar{y})^2}{k}$$

where $t$ is the frame number and $(x, y)$ are the coordinates of the centroid. Values of $V$ for unbound and plasmid-bound probes were higher than for stuck particles as across the whole length of the video, since they explored the whole pit due to random diffusion (SI Fig. 6).

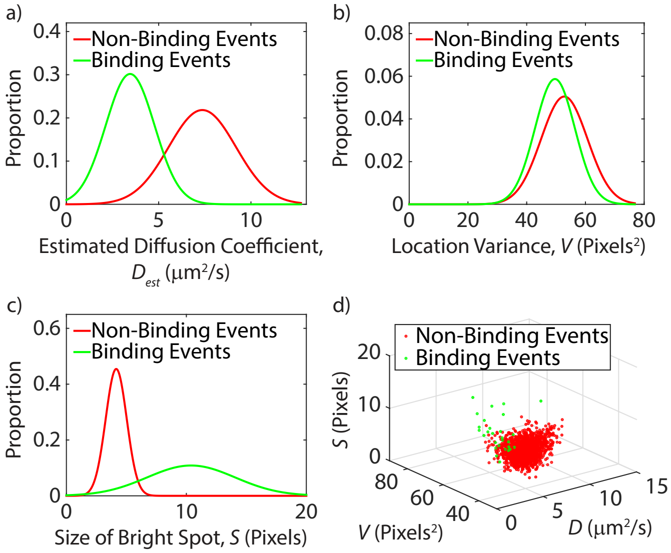


**Supplementary Figure 7: Predictors for a sample data set.** Predictors calculated for a sample data set of 90 videos (~36,000 data points), with binding events shown in green and non-binding events in red. **a)** The probability density curve for $D_{est}$ for this data set. **b)** The probability density curve for $V$ for this data set. **c)** The probability density curve for $S$ for this data set. **d)** Three-dimensional scatter plot of all predictors ($D_{est}$, $V$, and $S$), for a sample size of 3 videos (~1200 data points) from this data set.

#### 10.4 Calculating Bright Spot Size

Finding the precise outline of the above "spots" allowed us to track the diffusing plasmid-probe complexes. For each frame, the bright spot size $S$ was found by the flood-fill algorithm. For the most part, the plasmid-probe complexes had larger $S$ values since they appeared as a single concentrated spots with homogeneous intensities surrounding the brightest pixel. This homogeneity allowed for a large connected component to form. Freely diffusing probes had lower $S$ values due to the lower, heterogeneous intensities surrounding the brightest pixel. This prevented a large connected component from forming (SI Fig. 6).

Furthermore, measuring the distribution of $S$ allowed us to identify "outlier" probe aggregates, which tended to be larger (and brighter) than the probes and probe-plasmid complexes.

#### 10.5 Significance of Predictors

The distribution of each predictor (Estimated Diffusion Coefficient $D_{est}$, Size of Bright Spot $S$, and Location Variance $V$) of the plasmid-bound and unbound probe data sets were fitted to Gaussian distributions and plotted for a sample data set (SI Fig. 7). To quantitatively deter-

**Supplementary Table: Sample predictor values**

| Predictor | p-value | Significance |
|---|---|---|
| $D_{est}$ | $< 10^{-16}$ | Significant |
| $V$ | $5.35 \times 10^{-9}$ | Significant |
| $S$ | $< 10^{-16}$ | Significant |

mine whether the distributions of the plasmid-bound and unbound sets were significantly different for each predictor, the two-sample Kolmogorov-Smirnov test was used [5]. This statistical test is based on the null hypothesis that two distributions are drawn from the same parent distribution. The results for this test on a sample data set are shown in Table 1. Since all $p$-values were below $\alpha = 0.05$, the null hypothesis for each predictor was rejected. This means that the bound data set and unbound data set cannot be from the same parent set, or that they were found to be significantly different from one another.

#### 10.6 Constructing the K-NN Classifier to Detect Probe-plasmid Complexes

A distance-weighted K-nearest neighbor (K-NN) classifier was utilized to distinguish bound probes from unbound probes based on their predictors: $D_{est}$, $V$, and $S$. The output of the classifier was a Boolean variable, representing whether a certain pit contained a bound probe or not. A supervised machine learning approach with continuous training was used to improve accuracy while a data set was analyzed. First, a training set was constructed and continuously improved upon by the user with a custom GUI. This GUI displayed one of the videos for a sample data set, allowing the user to verify by eye which pits contained probe-plasmid complexes. A grid representing each pit in the video allowed the user to input which pits contained binding events in the video of interest. Applying the user inputs from the grid, the algorithm updated the predictors ($D_{est}$, $V$, and $S$) in the training data set, allowing more accurate automatic detection of probe-plasmid complexes. Thus, the user was able to correct any false positives or negatives that the algorithm generated, further improving the training set as more videos were analyzed. The K-NN algorithm was formally defined as:

**procedure** KNN(trainingSet $T$, query $x$)

Let $t_1..t_k \in T$ be the $k$ nearest neighbors of $x$.

Let $w_i = (D_{t_i} - D_x)^2 + (S_{t_i} - S_x)^2 + (V_{t_i} - V_x)^2$

Compute $Score(x) = \sum_{i=1}^{k} w_i^{-1} L(t_i)$

where $L(t_i) = \begin{cases} 1 & \text{if } t_i \text{ contains bound probe} \\ -1 & \text{otherwise} \end{cases}$

**if** $Score(x) > 0$ **then return** Bound

**else return** Not-Bound

Tenfold cross-validation tests were conducted in order to determine the optimal parameter, $k$ (number of neighbors). The cross-validated error (measuring classifier performance) was minimized at 10 neighbors, thus $k = 10$ was chosen.

#### 10.7 K-NN Classifer Performance

To quantify how accurate the K-NN classifier was in detecting probe-plasmid complexes, the sensitivity rate was calculated as:

$$\text{Sensitivity} = \frac{\text{TP}}{\text{P}} = 1 - \text{FNR} \quad (2)$$

where TP and P were the number of pits with probe-plasmid complexes identified by the algorithm and by the user, respectively. After the analysis of approximately 20 videos, a sensitivity of $\approx 75\%$ was achieved for detecting probe-plasmid complexes. This corresponded to a false negative rate of $\approx 25\%$.

Another quantitative measure of the K-NN classifier performance could be accomplished by considering the number of false positives made by the algorithm. This is quantified by the specificity rate, defined as:

$$\text{Specificity} = \frac{\text{TN}}{\text{N}} = 1 - \text{FPR} \quad (3)$$

where TN and N were the number of pits without probe-plasmid complexes identified by the algorithm and by the user, respectively. Throughout the analysis process, the specificity averaged around 99.99%, equivalent to a false positive rate of 0.01%.

The algorithm was trained and supervised throughout the analysis process. All of the results presented in this work were checked manually by two independent users, to correct for potential errors.

#### 10.8 Probe Counting

For accurate probe-plasmid binding statistics, it was necessary to count the total number of probes under consideration in a video, allowing for accurate scaling and comparison between different data sets. To do so, a probe counting algorithm was implemented.

As the illumination over an entire video was not uniform, some pits, usually around the edges, were too dark for robust probe counting. To eliminate dark pits from consideration, the flood-fill algorithm described above was ran twice. In contrast to the earlier algorithm, the input is the matrix $P$, where $P_{ij}$ was used to represent the mean intensity of the $ij^{th}$ pit. The seed was selected to be the brightest pit out of the entire pit array. Expansion occurred as described previously, but diagonal neighbors were also taken into account. For the first pass of the algorithm, the termination threshold was set to be the mean intensity of the darkest pit (empty pit) within the brightest 8x8 array of pits. This generated a connected component of bright pits, but with gaps. A second pass of the flood-fill algorithm was used, this time with the termination condition of having less then 3 neighbors which were explored in the first pass. This served to fill in gaps of the bright region. Pits outside of the resulting region were discounted from the counting algorithm.

Next, the total number of probes were counted by considering each pit (which was determined to be bright enough for robust consideration). The overall spatial variance in intensity for each pit and its surrounding background in a single video were obtained by averaging the spatial variance over each frame of the video. Formally, for a single frame, the spatial variance is defined as:

$$SpatVar = \sum_i \sum_j (I_{i,j} - Mean(I))^2 \quad (4)$$

For each video, the pit's "actual intensity variance" was calculated using the difference between the pit pixels and the surrounding background pixels, to correct for the autofluorescence of the glass. Using a data set which was obtained for a dilute sample, in which each pit contained either zero or one probe molecule, a scale factor was determined to convert the "actual intensity variance" to probe number. The probe number distributions were then binned (from 0 probes in a pit, to the maximum detected probe occupancy in a pit). This histogram was then fit to a Poisson distribution, and the mean was taken to represent the average number of probes per pit for a video.

##### 10.9 Spatial Variance in Intensity and Blinking

As fluorophores are also known to blink, a temporary state where it temporarily is incapable of emitting photons, it was possible that the increase in the spatial variance in intensity in the Section Experimental results: binding kinetics was due to a fluorophore which had been blinking. To control for this, we monitored the pit's median intensity over time. If a molecule was blinking, it would represent either an increase or decrease in the median pit intensity at the same time that we observed a shift in the spatial variance in intensity. For all binding events captured in our videos we observed no significant change in the median intensity of the pit, indicating that the fluorophore that was captured was emitting at all times and was not blinking. Thus, the change in the spatial variance in intensity must be due to a change in the diffusion of the fluorescent probe caused by binding to an unwound plasmid.

Using this technique, it was also theoretically possible to detect whether unwinding sites would close over the course of an experiment, resulting in probes unbinding. Over all of our experiments, no unbinding events were detected using this algorithm. Subsequent experiments where probes were incubated with plasmids at 37°C overnight under conditions where binding should occur demonstrate complete binding between probes and plasmids (see SI Section 8). From this, two conclusions were made. First, this indicates that equilibrium for this system occurs at time scales longer than our experiments. Secondly, this indicates that once a probe was bound, it remained bound to the plasmid during the course of an experiment. These support the idea that rewinding of the plasmids is either very slow, or becomes impractical with a bound probe.

#### 11 Guiding Predictions of SIDD and DZCBtrans Models

Theoretical calculations of strand unwinding were performed using both the SIDD algorithm and the DZCBtrans algorithm. Our comparison of model predictions with experimental results supports the importance of including the competition between higher order structural transitions, such as Z-form, cruciform, and strand denaturation, in studying this system.

##### 11.1 SIDD Model Predictions

Theoretical predictions presented in this work were first made using the SIDD algorithm, detailed in the work of Zhabinskaya & Benham [6, 2]. The SIDD algorithm predicts the locations and extent of stress-induced duplex destabilization (SIDD) in DNA in the absence of other transitions. The technique, which was described in the paper, was applied to calculate the extent of strand separation in pUC19 plasmids as a function of temperature and superhelical density. Samples used for the microscopy experiments contained a heterogeneous mixture of pUC19 at different superhelical densities, as described in the section Topoisomer Production. To generate theoretical predictions that accurately represent our topoisomer samples, the weighted mean of the number of open base pairs is calculated, weighted according to the amount of each topoisomer present. More precisely, for each sample with mean superhelical density $\langle\sigma\rangle$, and a distribution of topoisomers $\sigma_1, ...\sigma_n$, the fraction of each topoisomer within the sample, $w_j$, was found by gel analysis tools as described in the Topoisomer Production section. For each base pair, $x_i$, the weighted mean of denaturation probability ($\langle P\rangle$) for a sample with mean superhelical density $\langle\sigma\rangle$ was defined as:

$$\langle P(x_i)\rangle = \sum_{j=1}^{n} w_j P_{\sigma_j}(x_i) \quad (5)$$

where $P_{\sigma_j}$ was the denaturation probability at a specific superhelical density, $\sigma_j$. These weighted means were then summed over specific sequence regions to determine the predicted number of open base pairs in each region.

Theoretical predictions using only SIDD profiles without considering other structural transitions are given in SI Fig. 8. For these results, the ionic strength parameter was set to 22.5 mM, as determined by the buffer composition used during microscopy. Comparing this with Fig. 2 of the main text, it is evident that a much higher number of bases are destabilized at lower temperatures if only strand separation is considered. This is in contrast to our data, whose low relative binding at these conditions, points to the importance of taking other structural transitions into account.

##### 11.2 DZCBtrans Model Predictions

Since SIDD does not consider competition between strand unwinding and other kinds of secondary structures (such as Z-DNA and cruciforms), we employed the DZCBtrans algorithm developed by Zhabinskaya and Benham to determine what effects these structures would have on denaturation [2]. The DZCBtrans algorithm has been shown to accurately predict the local DNA structure by considering the base sequence to find regions that are susceptible to Z-form, cruciform, or strand denaturation. We note that certain sequences can be susceptible to more than one type of alternate conformation.

Z-DNA is a left-handed helix whose structure resembles a "zig-zag" pattern [7]. Z-DNA is known to prefer alternating purine-pyrimidine sequences, specifically $(GC)_n$ or $(CG)_n$ runs, but can also occur in other base sequences [2]. Negatively supercoiled plasmids favor Z-DNA at low temperatures as under these conditions other types of alternate structures are not competitive [8]. The predicted probabilities of Z-DNA formation at each base pair position of pUC19 at 31°C is shown in SI Fig. 9a as a function of $\sigma$. The overall predicted number of bases of pUC19 with Z-DNA structure are shown in SI Fig. 9b as a function of $T$ and $\sigma$. Significant Z-DNA formation is apparent at highly negative supercoiling levels and low temperatures, which tapers off at higher temperatures and lower overall supercoiling. Using the DZCBtrans algorithm, we observe no significant predicted cruciform formation in pUC19 for all temperatures and superhelical densities. Results for strand denaturation predicted by the DZCBtrans algorithm at the Site 1 region (bp 1525 - bp 1633) with Z-form and cruciform competition taken into account are shown in Fig. 2 of the main text.

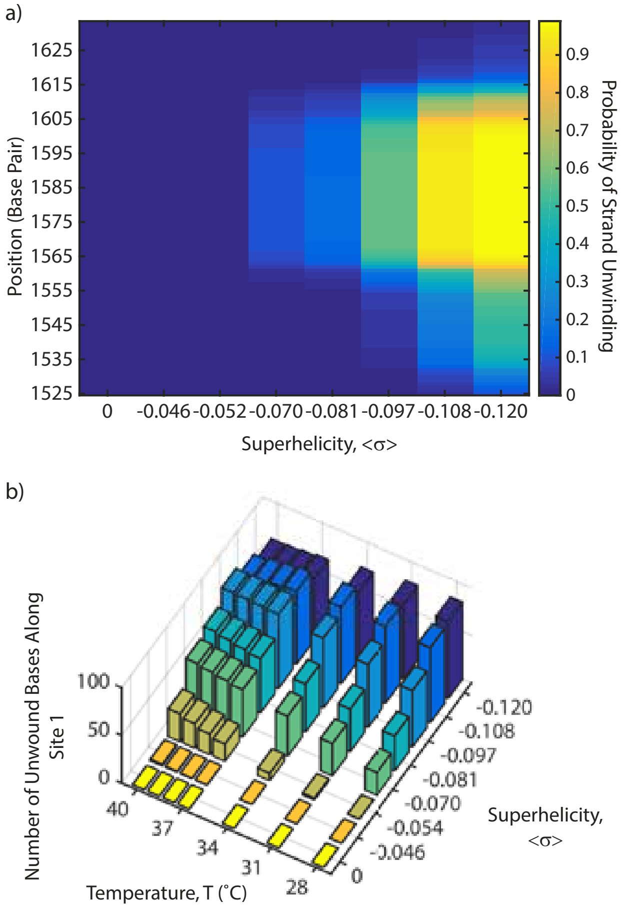


**Supplementary Figure 8: Stress Induced Duplex Destabilization (SIDD) of pUC19. a)** Probability map illustrating the locations within Site 1 of pUC19 undergoing SIDD, at 37°C, as a function of $\sigma$. **b)** Predicted number of bases undergoing SIDD vs. temperature and superhelicity in the Site 1 region of pUC19. This prediction only considers strand denaturation and does not account for local competition between strand-unwinding, B-DNA, Z-DNA, and cruciform formation (DZCBtrans). Note the high number of destabilized bases at low temperatures for higher superhelicities, in contrast to that of DZCBtrans in Fig. 2 and below. The discrepancy is due to B-Z transitions occurring at low temperatures. The SIDD behavior at average superhelicity, $\langle\sigma\rangle$ takes into account a Gaussian distribution of superhelical densities centered around $\langle\sigma\rangle$ which reflects the distribution found in our experimental topoisomer samples.

#### 12 Reaction Kinetics Model Fits

The fits presented in this work were derived from the second-order reaction equation. For unbound probes $O$

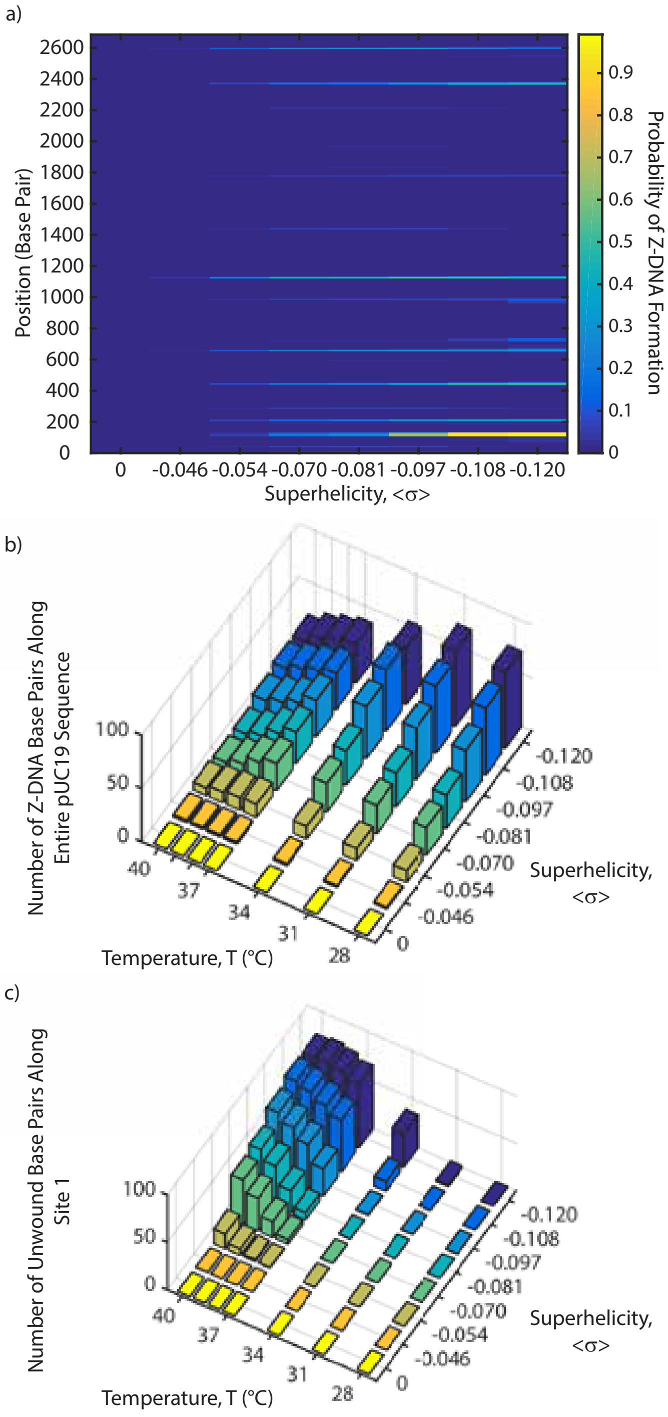


**Supplementary Figure 9: Z-DNA locations of pUC19.** **a)** Probability map illustrating the base pair locations of pUC19 with Z-DNA structure at 31°C as a function of $\sigma$. **b)** Predicted number of bases with Z-DNA structure across the entire pUC19 sequence. Significant Z-DNA formation occurs at highly negative supercoiling levels and at low temperatures. **c)** Predicted number of unwound bases across Site 1, calculated using the DZCB-trans algorithm for comparison purposes.

and unwound plasmids $U$, this reaction is defined as

$$\ln\left(\frac{[U][O]_0}{[O][U]_0}\right) = k\left([U]_0 - [O]_0\right)t \qquad (6)$$

where $k$ is the rate of reaction as defined in the text, and $t$ is the time passed since the reaction began.

In our model, we assumed that both the initial concentration of unwound plasmids $[U]_0$ and the interaction rate $k$ were free to vary with temperature. It is possible to create a model where either $k$ or the $[U]_0$ is held constant with temperature $T$. To determine which would fit our data best, we calculated the $\chi^2$ values of all three models ($k$ is constant with $T$, $[U]_0$ is constant with $T$, or both vary with $T$). $\chi^2$ is a measure of how well a mathematical model fits a data set, with the lowest value of $\chi^2$ indicating the best fit. From these values of $\chi^2$, we determined that the model that allows both $k$ and $[U]_0$ to vary with $T$ gave the best fit (data not shown), and so this model was selected to fit the data in this work.

#### 13 *E. coli* Topoisomerase IA Reaction Kinetics

21.1 nM of supercoiled pUC19 was mixed with 752 pM of the 30-base probe (described above) in a buffer of 20 mM tris, 50 mM potassium acetate, and 100 $\mu$g/mL BSA (final tris concentration was 18.42 mM). 25 $\mu$L of this solution was incubated at 37°C overnight for a total of 16 h prior to observation on the microscope.

Topoisomerase IA was purchased from from New England Biolabs (catalog # M0301L). Topoisomerase IA is distinguished from topoisomerase IB by the fact that the former relaxes only negative supercoils, while the latter can relax both positive and negative supercoils.

The reaction buffer used in the experiment was different from the ideal buffer for topoisomerase IA reactions, namely in that magnesium was not included as it caused DNA to stick to the flow cell. To test whether modifications of the NEC recipe affected the topoisomerase IA reaction, the reaction was performed in a PCR tube on supercoiled pUC19 in a range of modified reaction buffers. Topoisomerase IA was added to each sample, and then samples were incubated for 2 h at 37°C. The reaction was stopped by heating the samples to 65°C for 20 min, and the samples were stored at 4°C until use.

To confirm DNA supercoil relaxation, the same topoisomerase IA reaction as one performed on the microscope was run in bulk solution and run on a series of agarose gels (Fig. 10). As the microsocpy flow cells were pre-treated with 55 kDa PVP which was then exchanged with buffer, the actual amount of PVP in solution during the experiment was unknown. As such, the topoisomerase IA reactions were carried out in various concentrations of PVP. The reaction was carried out both with and without incubating the samples with probes to determine if the they

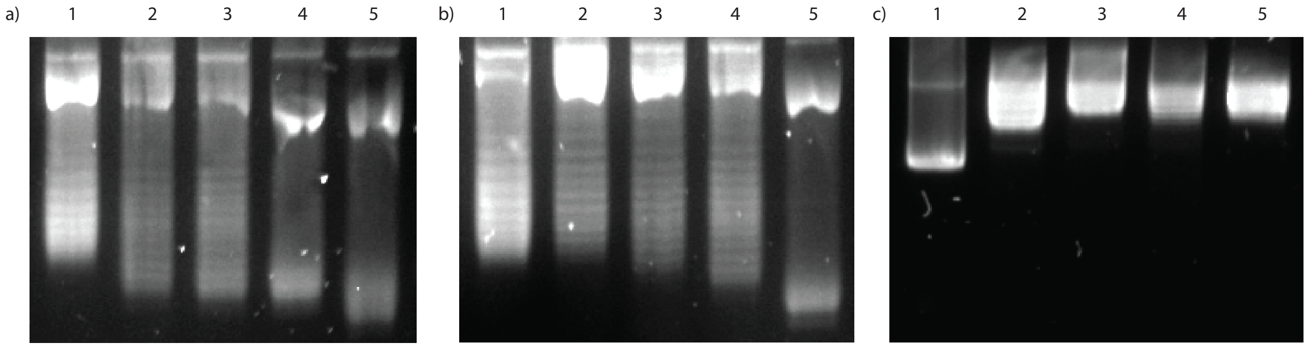


**Supplementary Figure 10: Series of agarose gels demonstrating DNA supercoil relaxation by topoisomerase IA. a)** 2% agarose gel run at 30 V in the presence of chloroquine diphosphate for 40 h. Lane 1: supercoiled pUC19. Lanes 2-4: supercoiled pUC19 treated with topoisomerase 1A with 1%, 0.1% and 0% 55 kDa PVP, respectively. Lane 5: supercoiled pUC19 treated with topoisomerase 1A in the commercial buffer. **b)** same as a), except 752 pM of fluorescently-labelled 30-base probes were added to samples 1-4, and incubated at 37°C overnight before the addition of topoisomerase.**c)** same as a) but run on a 1% agarose gel at 120 V for 1 h with no chloroquine diphosphate. All gels were run in a 1x TAE buffer composed of 40 mM Tris, 20 mM acetic acid, and 1 mM EDTA with 6 $\frac{\mu g}{mL}$ chloroquine diphosphate, except where indicated otherwise.

interfered with the topoisomerase activity (Fig. 10a) and b), respectively). Agarose gels were run as detailed in SI Section 9 in the presence of 6 $\frac{\mu g}{mL}$ of chloroquine diphosphate. A final agarose gel, using identical solutions to Fig. 10a), was run with 1% agarose for 1 h with no chloroquine diphosphate.

The supercoiled sample in Lane 1 should run a different distance in the gel from the supposedly relaxed DNA molecules in Lanes 2-5 in Fig. 10a) and b). As shown here, this was not the case. However, as the choloroquine in the gel may have positively supercoiled the DNA in Lanes 2-5 such that they ran similar distances on a gel, a 1 h gel was run at 120 V with no chloroquine diphosphate to see if they ran differently. From Fig. 10c), the unreacted DNA in Lane 1 ran much further than the topoisomerase-reacted DNA in any other lane. This would only occur in a non-chloroquine gel if it was much more supercoiled than the DNA in the other lanes, indicating that the topoisomerase reaction worked in the samples demonstrated on Lanes 2-5.

Topoisomerase IA activity is partially inhibited by the experimental buffer comprised of 50 mM potassium acetate, 20 mM Tris, 100 $\frac{\mu g}{mL}$ BSA with no PVP (Lane 4) and with the presence of PVP (Lanes 2-3). While the topoisomerase reaction didn't run to completion in the non-commercial buffer, for the purposes of this study it was sufficient to have sufficient supercoil relaxation to cause the probe to unbind.

A flow cell was prepared following the above procedure, using a 3-$\mu$m pit array embedded in the bottom coverslip. The flow cell was mounted onto the microscope. To further prevent the sample from sticking to the flow cell, 26 $\mu$L of 10% polyvinylpyrrolidone (PVP, Sigma-Aldrich Canada Co., Oakville, ON, catalog # PVP10-100G) in 90 mM tris was mixed with 2.5 mM of PCA and 50 nM of PCD and inserted into the flow cell. After 10 min, the PVP solution was rinsed out of the flow cell with a buffer solution which was characterized by 26 $\mu$L of 20 mM tris, 50 mM potassium acetate, 100 $\mu$g/mL BSA, 2.5 mM of PCA, and 50 nM of PCD. The temperature control on the microscope was set to 37°C and the flow cell was exposed to green (532 nm) laser light for 10 min to photobleach any impurities in the glass and objective oil.

Prior to pipetting the samples into the CLiC device, PCA, PCD, and *E. coli* topoisomerase IA (New England Biolabs, Whitby, ON, catalog # M0301L) were added for final concentrations of 2.5 mM, 50 nM, and 28.48 $\mu$g/mL respectively. The sample was inserted into the device and a 50-frame video was taken every minute for two hours after the enzyme was added. The sample was mixed between videos by raising and lowering the CLiC lens. The first video was taken 290 s after the enzyme was added to the sample. The amount of binding was analyzed for three videos every 15 min using the binding detection algorithms described above. As described in the main text, the probes were observed to unbind from the plasmids over time.
